# Cooperative Binding of Transcription Factors is a Hallmark of Active Enhancers

**DOI:** 10.1101/2020.08.17.253146

**Authors:** Satyanarayan Rao, Kami Ahmad, Srinivas Ramachandran

**Affiliations:** Department of Biochemistry and Molecular Genetics, Fred Hutchinson Cancer Research Center, 1100 Fairview Avenue, North Seattle, WA 98109, USA; RNA Bioscience Initiative, University of Colorado School of Medicine, Mail Stop: 8101, 12801 East 17th Ave. L-18-9102, Aurora, CO 80045; Basic Sciences Division, Fred Hutchinson Cancer Research Center, 1100 Fairview Avenue, North Seattle, WA 98109, USA

## Abstract

Enhancers harbor binding motifs that recruit transcription factors (TFs) for gene activation. While cooperative binding of TFs at enhancers is known to be critical for transcriptional activation of a handful of developmental enhancers, the extent TF cooperativity genome-wide is unknown. Here, we couple high-resolution nuclease footprinting with single-molecule methylation profiling to characterize TF cooperativity at active enhancers in the *Drosophila* genome. Enrichment of short MNase-protected DNA segments indicates that the majority of enhancers harbor two or more TF binding sites, and we uncover protected fragments that correspond to co-bound sites in thousands of enhancers. We integrate MNase-seq, methylation accessibility profiling, and CUT&RUN chromatin profiling as a comprehensive strategy to characterize co-binding of the Trithorax-like (TRL) DNA binding protein and multiple other TFs and identify states where an enhancer is bound by no TF, by either single factor, by multiple factors, or where binding sites are occluded by nucleosomes. From the analysis of co-binding, we find that cooperativity dominates TF binding *in vivo* at a majority of active enhancers. TF cooperativity can occur without apparent protein-protein interactions and provides a mechanism to effectively clear nucleosomes and promote enhancer function.

## Introduction

Cis-regulatory elements (CREs) or enhancers are DNA sequences that drive cell type-specific gene expression, developmental transitions, and cellular responses to external stimuli (Banerji et al., 1981; Dunipace et al., 2011; Lagha et al., 2012; Levine, 2010). In eukaryotes, CREs are usually ~500 bp in length with multiple binding sites for transcription factors (TFs). A fundamental question in gene regulation is: what is the role of multiple TF binding sites in driving enhancer function? First, multiple binding sites could increase genomic specificity of CREs, as single transcription factor binding sites (TFBS) are short and thus occur often by chance in large genomes but multiple juxtaposed TF sites are less likely (Crocker et al., 2015; Ludwig et al., 2011). Second, multiple sites provide higher affinities than individual motifs (von Hippel and Berg, 1986). Third, multiple TFBSs at enhancers may be required to program cell type specificity with combinations of TFs (Lagha et al., 2012).

Additional reasons for juxtaposing multiple factor binding sites arise from considering that enhancers must function in chromatin. In the genomes of multicellular organisms, most enhancers are occluded by nucleosomes when not active (Schones et al., 2008), arguing that TF binding and not just the underlying sequence features are required to expose the DNA in the regulatory element. In a hierarchical model of enhancer function, binding of one initiating TF may displace nucleosomes and expose binding sites for other secondary TFs (Iwafuchi-Doi and Zaret, 2014). Alternatively, in a billboard model, multiple TFs may independently bind to an enhancer, and any one TF may keep the regulatory element nucleosome-free (Arnosti and Kulkarni, 2005; Reiter et al., 2017; Spitz and Furlong, 2012). Finally, in an enhanceosome model, protein-protein interactions between bound TFs may drive nucleosome displacement and enhancer function (Bintu et al., 2005; Joshi et al., 2007; Mann and Affolter, 1998). TF occupancy at enhancers differentiates these models for enhancer function; in hierarchical and billboard models, an initiating TF might spend more time bound at an enhancer, but with little co-binding with other factors. In contrast, in enhanceosome-like complexes, co-bound states will be frequently observed. To characterize TF occupancies across a genome, we need to be able to map TF binding at high resolution to distinguish independent and co-bound TFs.

Massively parallel reporter assays have now mapped the locations of thousands of enhancers in the genomes of defined cell types (Andersson and Sandelin, 2020; Arnold et al., 2013), setting the stage to characterize general rules for TF binding in regulatory elements. While traditional methods such as Chromatin Immuno Precipitation (ChIP) for mapping bound transcription factors have poor resolution and sensitivity, more recent chromatin profiling methods such as ChIP-exo (He et al., 2015; Rhee and Pugh, 2011), ORGANIC native-ChIP (Kasinathan et al 2014) and CUT&RUN (Skene and Henikoff, 2017) now provide base-pair resolution. Additionally, DNase- and MNase-based methods can also be used to map TF footprints *in vivo*. While partial digestion with the endonuclease DNase primarily measures accessibility, limit digestion with the endo-exonuclease MNase produces DNA fragments protected from digestion by bound chromatin proteins (Henikoff et al., 2011; Hesselberth et al., 2009). These methods can be used to infer the accessibility and factor binding genome-wide.

Nuclease-based methods chew apart chromatin particles, losing information of what particle co-existed on a single chromatin strand. In contrast, mapping protein binding with exogenous DNA methyltransferases preserves DNA molecules and information on neighboring particles. One such method is dual-enzyme Single-Molecule Footprinting (dSMF) (Krebs et al., 2017), which uses both GpC and CpG DNA methyltransferases to methylate exposed DNA *in vivo*, thus identifying the positions of nucleosomes and bound transcription factors on a single molecule. The dSMF method also reveals states where neither TFs nor nucleosomes are bound, thus fully defining the occupancies of regulatory elements in the genome. Here, we combine high-resolution MNase-seq, ORGANIC ChIP, CUT&RUN, and dSMF to define TF binding events at enhancers in *Drosophila* S2 cells. We develop a method to map multiple TF binding at the same time using MNase-seq and CUT&RUN, inferring TF co-binding events. We measure the unbound state of an enhancer using dSMF, which enabled us to calculate cooperativity between co-binding TFs at enhancers. We find that co-binding is inversely related to nucleosome occupancy and stability, supporting models where TF cooperativity drives nucleosome displacement at active enhancers. Finally, the low occupancies of transcription factor binding sites in the *Drosophila* genome imply that transient TF binding and slow replacement of nucleosomes drives enhancer function.

## Results

### Active enhancers are enriched for short protected fragments

Limit treatment of chromatin with micrococcal nuclease (MNase) digests exposed DNA, producing fragments protected by histone octamers and by bound chromatin proteins (Henikoff et al., 2011). While histone-protected DNA is typically ~150 bp representing the length of DNA wrapped around a nucleosome, chromatin-bound transcription factors protect the DNA underneath their binding domains in the range of 10-50 bp lengths. Thus, short protected fragments should report TF binding at regulatory elements throughout the genome. To examine TF binding at enhancers, we used *Drosophila* S2 cells, where thousands of enhancers have been functionally mapped by STARR-seq (Arnold et al., 2013), a massively parallel reporter assay. STARR-seq reports DNA segments that promote transcription on a transient plasmid, and some of the recovered sequences are not active in the endogenous chromosomal location. The set of putative active enhancers is defined as DNase-hypersensitive STARR-seq sites (Arnold et al., 2013). Active and closed STARR-seq enhancers are also distinguished by active histone modifications such as H3K27ac and silencing modifications such as H3K27me3 (Arnold et al., 2013) (**Figure 1A, B**).

**Figure 1.**
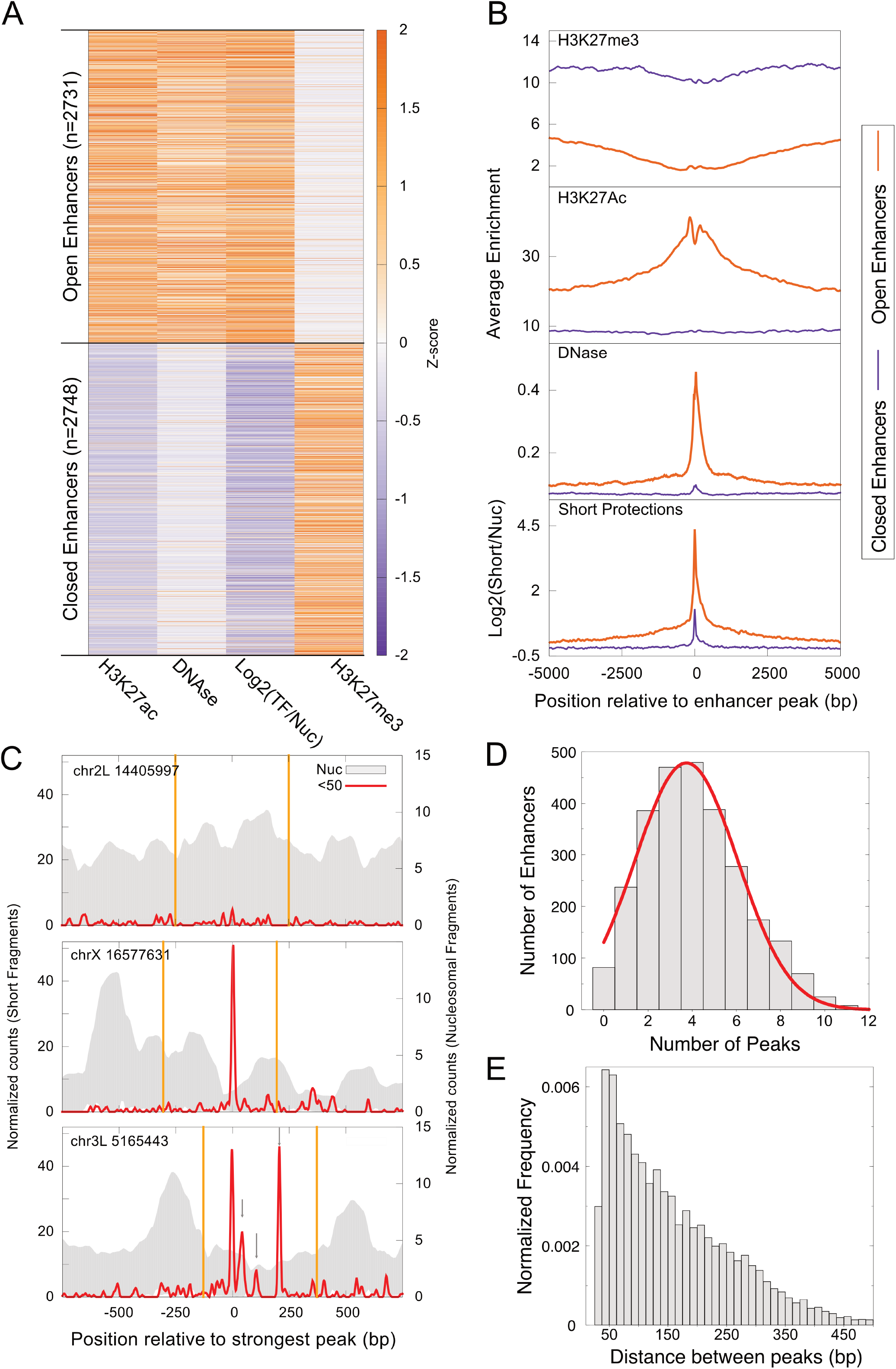
Active enhancers are enriched for short MNase-protected DNA fragments. **A**) Genome-wide enhancers (500 bp) in *Drosophila melanogaster* S2 cells identified by STARR-seq were classified into two clusters, active (n = 2,731) and closed (2,748) based on chromatin modifications (H3K27ac and H3K27me3 ChIP-seq data), DNase I hypersensitivity and enrichment (Log_2_) of MNase short (<50 bp) over nucleosomal (134-160 bp) protections. **B**) Enrichment of chromatin features at active and closed enhancers plotted relative to the primary MNase short protection peak. **C**) Examples from closed enhancer without any short MNase peak (top) and active enhancers demonstrating single (middle) and multiple (bottom) short MNase peaks. The gray background represents nucleosome occupancy. Peaks of the short fragment profile are indicated by gray arrows. **D**) Distribution of the number of MNase short protection peaks found at enhancers plotted with gray bars with a red line depicting the Gaussian fit. **E**) Distribution of distance between short MNase-protected peaks.

To determine if TF-protected fragments are detectable in active enhancers, we used sequencing data that is enriched for small fragments <100 bp by gel-isolation of MNase-digested chromatin (Ramachandran et al., 2017). We calculated a “short fragment” score as the log_2_ enrichment of DNA fragments <50 bp compared to nucleosome-sized fragments at 500 bp segments defined as enhancers in S2 cells by STARR-seq. We find that active enhancers are dramatically enriched for short protected fragments, while closed enhancers are depleted (**Figure 1A, B**). We conclude that active enhancers are abundantly occupied by short protected fragments, consistent with the binding of transcription factors in these regulatory elements.

### Many active enhancers contain multiple bound TFs

Visual inspection of individual STARR-seq sites confirms that active enhancers have specific short segments protected from MNase digestion, while these are absent from closed enhancers (**Figure 1C**). To map these putative TF binding sites, we called peaks on MNase-seq data in the <50 bp range. We used two stringent criteria to identify peaks: first, a peak must be a local maximum (>4 standard deviations from the mean); second, a peak must have >4-fold enrichment of short fragments over nucleosome-sized fragments. These criteria exclude nucleosomal intermediates at enhancer sites (Ramachandran et al., 2017). We observed that 96% of active enhancers have at least one peak. There is a wide number of peaks within enhancers: some enhancers have only one peak (**Figure 1C**, **middle**, **Figure 1D**), while others have clusters of multiple peaks (**Figure 1C**, **bottom**; **Figure 1D**). On average an active enhancer has four peaks (**Figure 1D**, Normal distribution: μ = 3.7, σ = 2.3), and in some cases, up to 11 peaks are distinguishable. The spacing between peaks within enhancers is very variable, with ~50 bp as the most observed distance (**Figure 1E**). These results suggest that small protected fragments can be used to report the configuration of TF binding events at high resolution at most active enhancers.

If short fragments within enhancers are protected from digestion by bound transcription factors, they should contain consensus motifs for those factors. We first scanned the short peaks within enhancers with collections of *Drosophila* transcription factor motifs and filtered matches for those factors that are expressed in S2 cells. The detected protected motifs are listed in **Supplementary Table S1**. One of the most abundant transcription factors in *Drosophila* S2 cells is the Trithoraxlike (TRL) protein, and indeed the consensus motif for TRL is strongly enriched within 36% of protected enhancer sequences that have a detected motif. To determine the correspondence of TRL binding to protection from MNase, we plotted the MNase short fragment enrichment and TRL mapping by ORGANIC native ChIP for active enhancers whose major protected peak displayed high-quality TRL motifs (**Figure 2A**). We observe a striking high-resolution correspondence between protected short fragments and bound TRL protein within enhancers. In contrast, enhancers without significant TRL motifs show no detectable TRL binding (**Supplementary Figure S1A**), and short fragments within these enhancers must be due to other transcription factors. Notably, aligning active enhancers by peaks of small protected fragments can resolve chromatin structural features within them.

**Figure 2.**
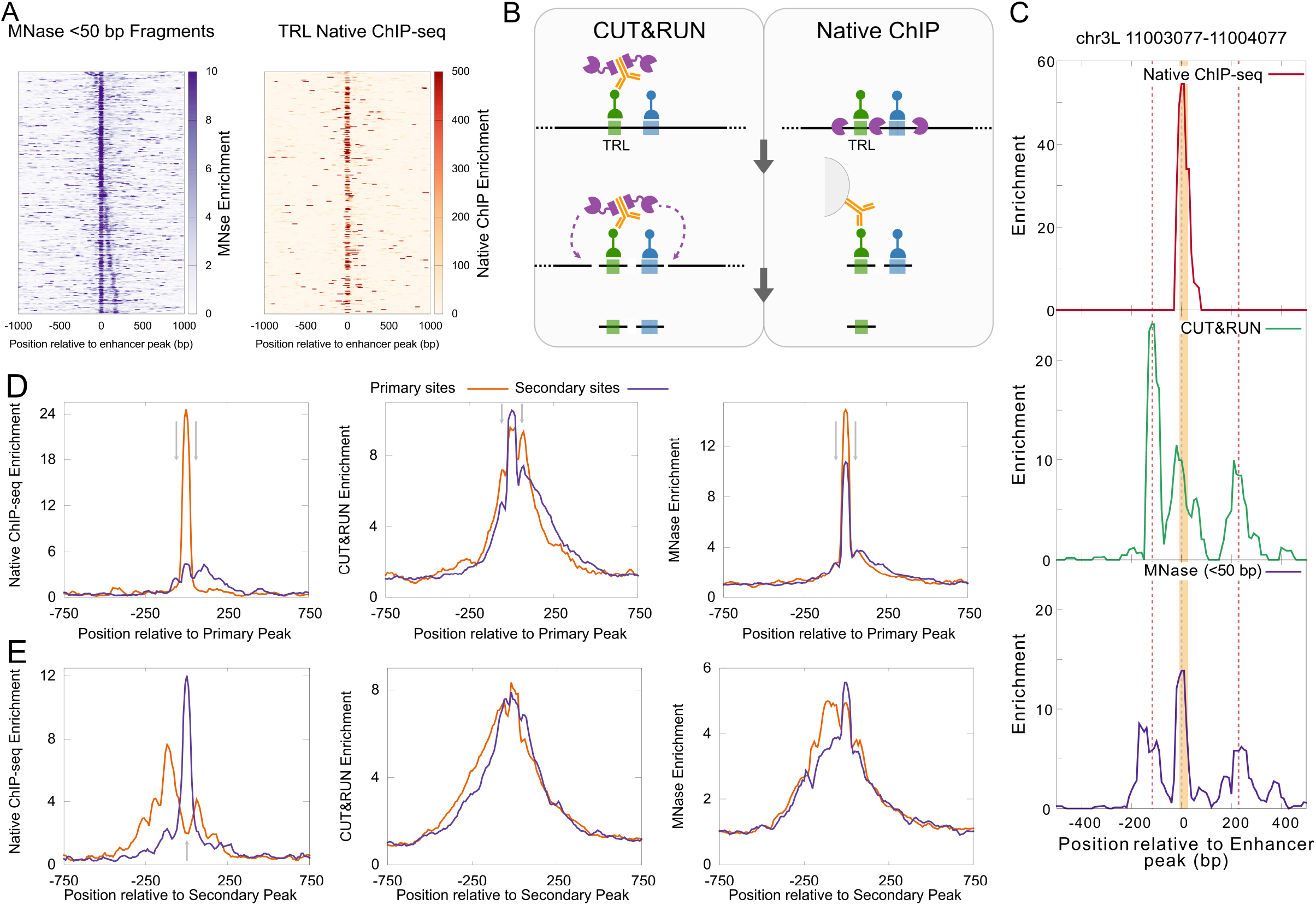
TRL binding at active enhancers. **A**) Heatmaps comparing enrichment of MNase short fragments and TRL ORGANIC native ChIP plotted relative to the primary peak of MNase short protection at enhancers with TRL motif at the central peak. **B**) Schematic demonstrating differences between CUT&RUN and ORGANIC in mapping co-bound TFs. The btlack line represents DNA and colored boxes on DNA represent motifs with TFs bound to them (green, TRL; blue, another TF). In ORGANIC (right panel) DNA is first treated with MNase (purple) before pull-down with an antibody (in orange), this leads to the loss of neighboring bound TFs. In CUT&RUN, protein A-MNase is tethered to a chromatin-bound antibody, and cleavage releases protein–DNA complexes in the vicinity of antibody-bound TRL. **C**) At a representative TRL binding site in an enhancer with no other TRL motifs, enrichment of TRL native ChIP-seq (top), CUT&RUN (middle), and <50 bp MNase-seq (bottom) are plotted. The gray dashed line represents the summit of the major Mnase peak. The red dashed lines represent the secondary peaks. TRL motifs overlapping peaks are shown as an orange shaded box. **D**) Enrichments plotted at enhancers centered at the primary MNase peak. These enhancers have single TRL motifs at either primary peak (Primary sites) or secondary peak (Secondary sites). ORGANIC short fragment enrichment (left), CUT&RUN short fragment enrichment (middle), and MNase short fragment enrichment (right) are plotted. Gray arrows at +/−60 bp depict enrichment of secondary sites which is strongest for CUT&RUN and weakest for ORGANIC. **E**) Same as (**D**) but the plots are centered at the secondary peak of enhancers with single TRL motif. Gray arrow in the ORGANIC plot is at the center and points to the depletion of signal for primary sites.

Immunoprecipitation methods like ORGANIC recover only the minimal fragment that is protected from MNase by a transcription factor or it’s protein complex because MNase-seq nibbles down all protected particles in a regulatory element (**Figure 2B**). However, profiling by CUT&RUN has the potential to preserve information from single DNA molecules. CUT&RUN uses an antibody to a chromatin protein to locally tether a protein A-MNase fusion, which then cleaves exposed DNA between proteins decorating that location (**Figure 2B**). Thus, in CUT&RUN data, any protected footprints around a factor binding site only appear if two factors are present on the same DNA molecule, for example, nucleosomes around chromatin-bound CTCF (Skene and Henikoff, 2017). We reasoned that we can extract information on the co-binding of two transcription factors by comparing ORGANIC, MNase-seq, and CUT&RUN profiles. Indeed, on an individual active enhancer, one main peak is detected in anti-TRL ORGANIC, and this peak precisely coincides with a high-quality TRL motif (**Figure 2C**). However, secondary peaks also appear on either side of the TRL-bound site in MNase-seq and in CUT&RUN data (**Figure 2C**). These secondary peaks in MNase-seq data can be either due to co-bound factors or independent binding of individual factors in the regulatory element, but in CUT&RUN profiling, the secondary peaks must be due to other transcription factors that are co-bound with TRL at the central peak.

We next compared ORGANIC, CUT&RUN, and MNase-seq across active enhancers with one TRL motif. If this co-binding happened without any protein-protein interactions, we would then observe high enrichment by ORGANIC at peak only when a high-quality TRL motif was present at that peak (**Supplementary Figure S1B**). Indeed, only motif-bearing primary sites show enrichment for TRL by ORGANIC at primary peaks, while secondary sites show background counts (**Figure 2D**, **left**). In contrast, we observed high enrichment for both primary sites and secondary sites when we plotted MNase-seq and TRL CUT&RUN at primary peaks (**Figure 2D**, **middle and right**). This pattern was reversed when we plotted TRL ORGANIC at secondary peaks. At secondary peaks, secondary sites showed high enrichment in ORGANIC, whereas primary sites showed a clear dip at the center with flanking peaks (**Figure 2E**, **left**). These results imply that co-bound transcription factors are common in these regulatory elements. Notably, secondary peaks are significantly higher in CUT&RUN profiling than in MNase-seq data, implying that factor-binding sites are often co-bound.

### High-resolution dissection of co-bound transcription factors

To identify the dominant combinations of multi-TF binding, we centered active enhancers at the main protected short fragment peak, and then performed k-means clustering (k=9) (**Figure 3A**). The resulting clusters revealed that ~28% of enhancers contain only one major peak of protected fragments, representing very simple enhancers (Cluster 3). In all other clusters, a secondary peak is also prominent and additional weaker peaks are also present. These are more complex multi-factor regulatory elements. Most enhancer clusters are depleted for nucleosomes across the region occupied by small fragments, thus corresponding to nucleosome-depleted regions (NDRs) that range in size from ~200 bp (Cluster 2) to as large as ~400 bp (Cluster 9) (**Figure 3B**). Cluster 1 is the exception: nucleosome depletion in this cluster is relatively weak, and this cluster also has the weakest small fragment peaks. Thus, the size and magnitude of NDRs are related to the spacing between small fragment peaks within the NDR, consistent with the antagonism between factors and histones for DNA.

**Figure 3.**
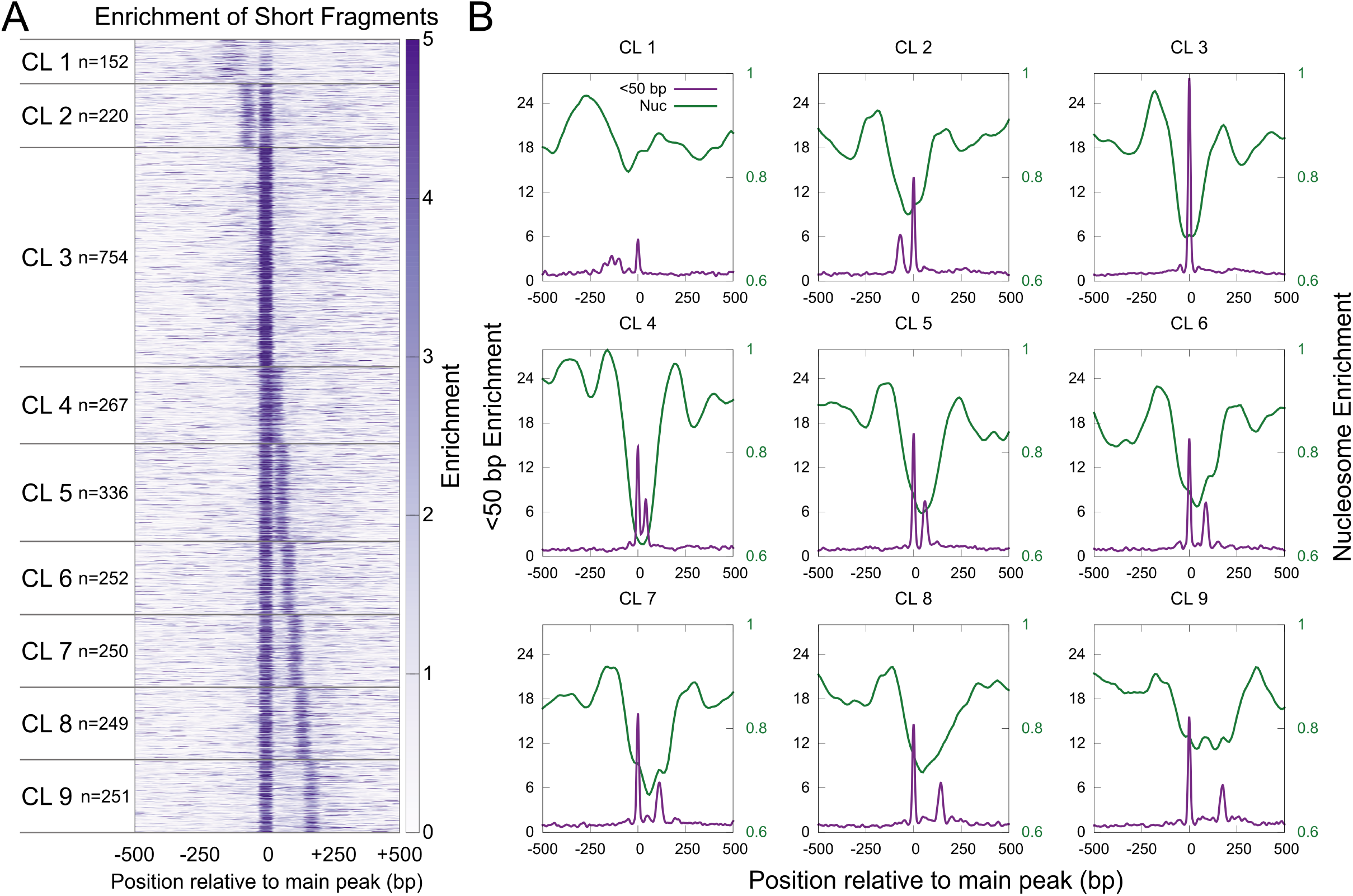
Binding of multiple transcription factors is common at enhancers. **A**) Heatmap of enrichment of short MNase-protected fragments plotted relative to the primary peak of short protection. Enhancers are ordered based on clusters defined by k-means. **B**) Average enrichment of short (<50 bp) and nucleosomal (134-160 bp) fragments of each enhancer cluster plotted relative to the primary peak of short protection.

To profile factor binding combinations in enhancers with two major peaks, we turned to V-plots. V-plots depict the density of DNA fragments as a function of their midpoint (x-axis) and their length (y-axis) (Henikoff et al., 2011). A chromatin-bound protein protects its minimal bound DNA from MNase digestion, but incomplete digestion on either end of the particle results in a notable “V” shape of plots, where the right line of the “V” arises from protection on the left side of the chromatin-bound protein, and the left line arises from protection on the right side. When aligned to the main peak, we observed strong “V” with the vertex at the peak center, pointing to minimal protection of ~40 bp for all the enhancer clusters, further confirming that we are mapping TF binding at enhancers (**Figure 4**, **Supplementary Figure S2**).

**Figure 4.**
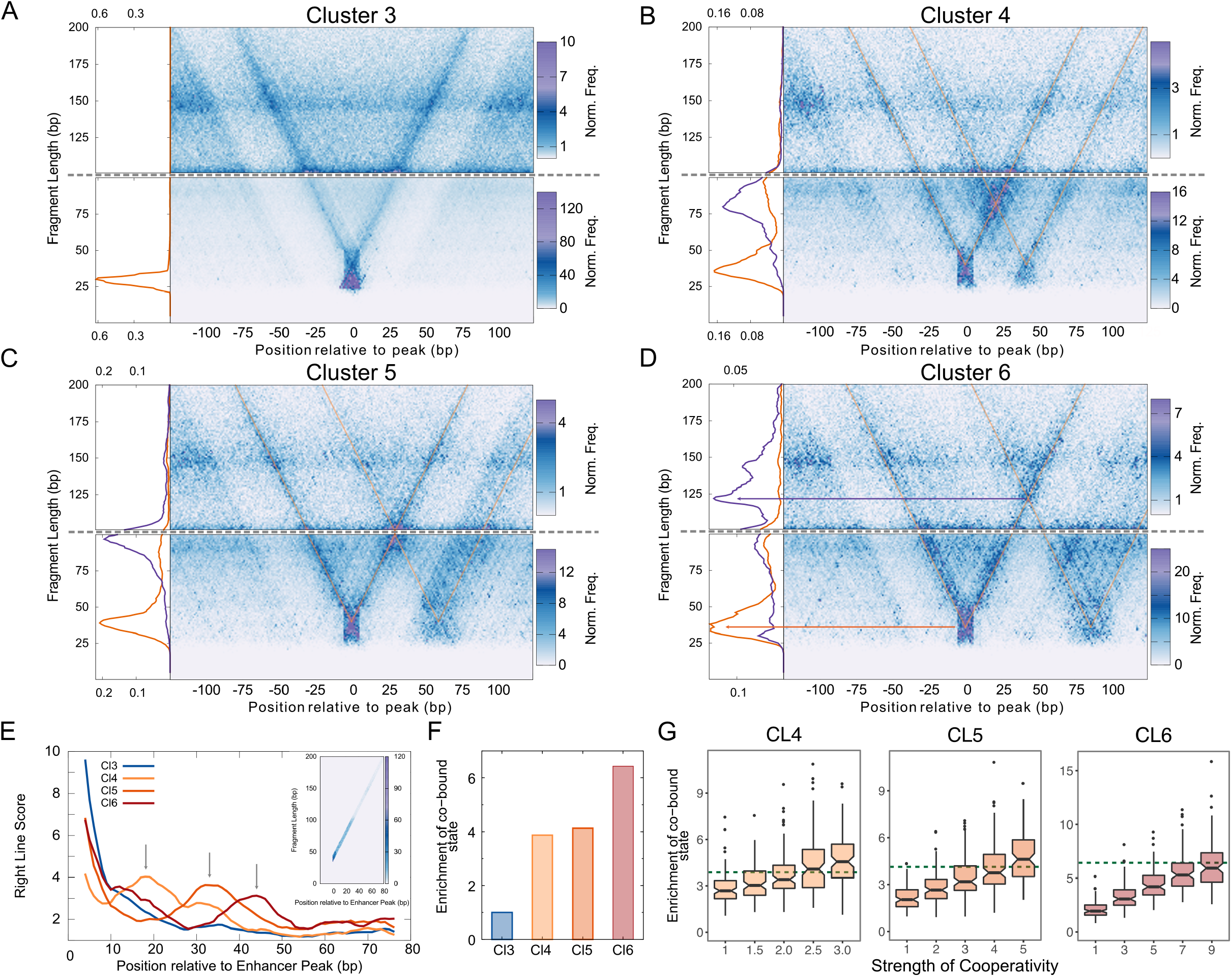
V-plot analysis reveals cooperative binding at active enhancers. **A**) Fragment midpoint versus fragment length plot (V-plot) centered at the primary peak for enhancers in Cluster 3 for fragments 100-200 bp (top) and 0-100 bp (bottom) are plotted separately. The average density of fragment lengths for fragments that map within 0±2 bp of the central peak is plotted to the left of the V-plots. **B**) Similar to (**A**) for Cluster 4. The average density of fragment lengths for fragments that map within 0±2 bp of the central peak (orange) and 21±2 bp of the central peak (the center of the co-bound species, purple) is plotted to the left of the V-plots. **C**) Similar to (**A**) for Cluster 5. The average density of fragment lengths for fragments that map within 0±2 bp of the central peak (orange) and 30±2 bp of the central peak (the center of the co-bound species, purple) is plotted to the left of the V-plots. **D**) Similar to (**A**) for Cluster 6. The average density of fragment lengths for fragments that map within 0±2 bp of the central peak (orange) and 43±2 bp of the central peak (the center of the co-bound species, purple) is plotted to the left of the V-plots. **E**) X-Z projection of the right line of the “V” from V-plot plotted for Clusters 3-6. The slice of the V-plot used in the projection shown in the inset. **F**) Enrichment of the co-bound states for Cluster 3-6 normalized to Cluster 3. **G**) Distribution of the enrichment of co-bound states for cluster 4 (left), 5 (middle), and 6 (right) normalized Cluster 3 from 100 simulated V-plots at each level of cooperativity indicated on the x-axis. The dashed line in each plot represents the observed enrichment from (**F**) for each cluster.

A V-plot of Cluster 3 enhancers centered on the primary small fragment peak displays a strong V with clear minimal protection of ~40 bp at the vertex, indicating the footprint of a single bound factor (**Figure 4A**). This vertex precisely corresponds to the peak of small protected fragments (**Figure 3B**, **Supplementary Figure S3**). In contrast, a V-plot for Cluster 4 enhancers shows three vertices (**Figure 4B**). Two vertices are minimal ~40 bp protected segments that correspond to the primary and secondary small fragment peaks of this cluster (**Figure 3B**, **Supplementary Figure S3**). The third vertex lies between the primary and secondary peaks with a fragment length of ~80 bp (**Supplementary Figure S3**). This position and length are consistent with a DNA fragment co-bound by TFs at both the primary and secondary sites. Other groups of enhancers also show the predicted arrangement of multiply bound TFs. Clusters 5 and 6 show a third vertex that always lies between peaks but increases size as these peaks are further apart (**Figures 4C, D**). For Cluster 5, the third vertex protects 100 bp, positioned 30 bp from the primary peak. For Cluster 6, the third vertex representing the co-bound particle is ~120 bp in size and is positioned 43 bp from the primary peak (**Supplementary Figure S3**). In Cluster 7-9, the size of the co-bound species approaches that of nucleosomes and is not clearly observed (**Supplementary Figure S2**).

From these V-plots, it is clear that the third vertex is formed by the right line of the primary peak V and the left line of the secondary peak V. This is because the left edge of the co-bound particle is the same as that of the TF bound at the primary peak, and the right edge of the co-bound species is the same as that of the TF bound at the secondary peak. Thus, the vertex of the co-bound particle will always lie on the right line of the primary peak V, and co-bound particles can be identified just by plotting the projection of the right line of the V (**Figure 4E**). For Cluster 3 there is a rapid decline of count density moving away from the sole peak (**Figure 4E**). Strikingly, for the multi-peak Clusters 4, 5, and 6, local accumulations are apparent at around 20, 30, and 40 bp respectively, indicating the positions of the co-bound vertex for each of those clusters (**Figure 4E**). The projection of the right line of the V confirms the observation of co-bound species in these clusters.

To estimate the abundance of co-bound TF particles, we calculated the ratio of fragment density at the co-bound vertex to the sum of fragment densities at the primary and secondary peak vertices. To account for differential recovery of short and long DNA fragments, these ratios are normalized by the ratio of counts at the same position in V-plots for Cluster 3, which lacks binding of a second TF. We define the strength of cooperativity as the ratio of the probability of observing co-bound state to the expected probability of co-binding if the two TFs were binding independently. We found a >3-fold excess of protection at the co-bound vertex compared to expectations from the binding of each TF (**Figure 4F**). To estimate the extent of cooperativity that would result in the observed enrichment of co-bound states, we simulated V-plots for different amounts of cooperativity. Strikingly, our simulations suggest the observed enrichment of co-bound states corresponds to ~2.5-fold excess of co-bound TFs compared to independent binding in Cluster 4, a ~4-fold excess in Cluster 5, and a ~8-fold excess for Cluster 6 (**Figure 4G**). Thus, many active enhancers show widespread and substantial cooperative binding of TFs.

### Identification of chromatin structure at enhancers

V-plots enable the identification of co-bound TFs at aggregated enhancers but cannot determine TF co-binding at sites on a single enhancer. Further, the fraction of factor binding sites that are not bound remains undefined. We turned to dual-enzyme Single-Molecule Footprinting (dSMF) (Krebs et al., 2017) to define binding states of individual enhancers. The *Drosophila* genome is devoid of cytosine methylation; thus, dSMF with exogenous GpC and CpG methyltransferases have been used to footprint chromatin proteins genome-wide. Critically, the dSMF method can capture information on multiple sites on a single DNA strand, allowing interrogation of all states of factor binding sites in an enhancer, including exposed sites, TF-bound sites, and nucleosome-occluded sites.

Active enhancers are accessible, while the DNA of closed enhancers is relatively inaccessible (**Figure 1A**). As expected, methylation by GpC and CpG DNA methyltransferases in the dSMF method depends on DNA accessibility and is starkly different between active and closed enhancers (**Figure 5A**). This suggests that factor-bound sites in active enhancers may be well-footprinted by the high density of DNA methylation in active regulatory elements. Therefore, we classified dSMF reads as “exposed” if all cytosines around a peak of small-fragment MNase-seq were methylated (**Figure 5B**, **top**). One or more contiguous unmethylated cytosines flanked by methylated cytosines in a dSMF read then defines protein-bound footprints on the enhancer, and we distinguished nucleosomes from TF-bound sites by the length of the unmethylated sequence, respectively (**Figure 5B**, **middle and bottom**; see ‘Assigning binding states to single factor sites in dSMF reads’ in Methods). Active enhancers are enriched for short unmethylated footprints, while closed enhancers are enriched for nucleosome-sized unmethylated footprints (**Figure 5E**). Footprint calls are not biased by sequence composition at enhancers, as we see no difference in length distribution of theoretical footprints (see ‘DNA molecule preparation and footprint calls’ in Methods) between active and closed enhancers (**Figure 5E**), confirming that dSMF footprinting can distinguish chromatin structures of regulatory elements.

**Figure 5.**
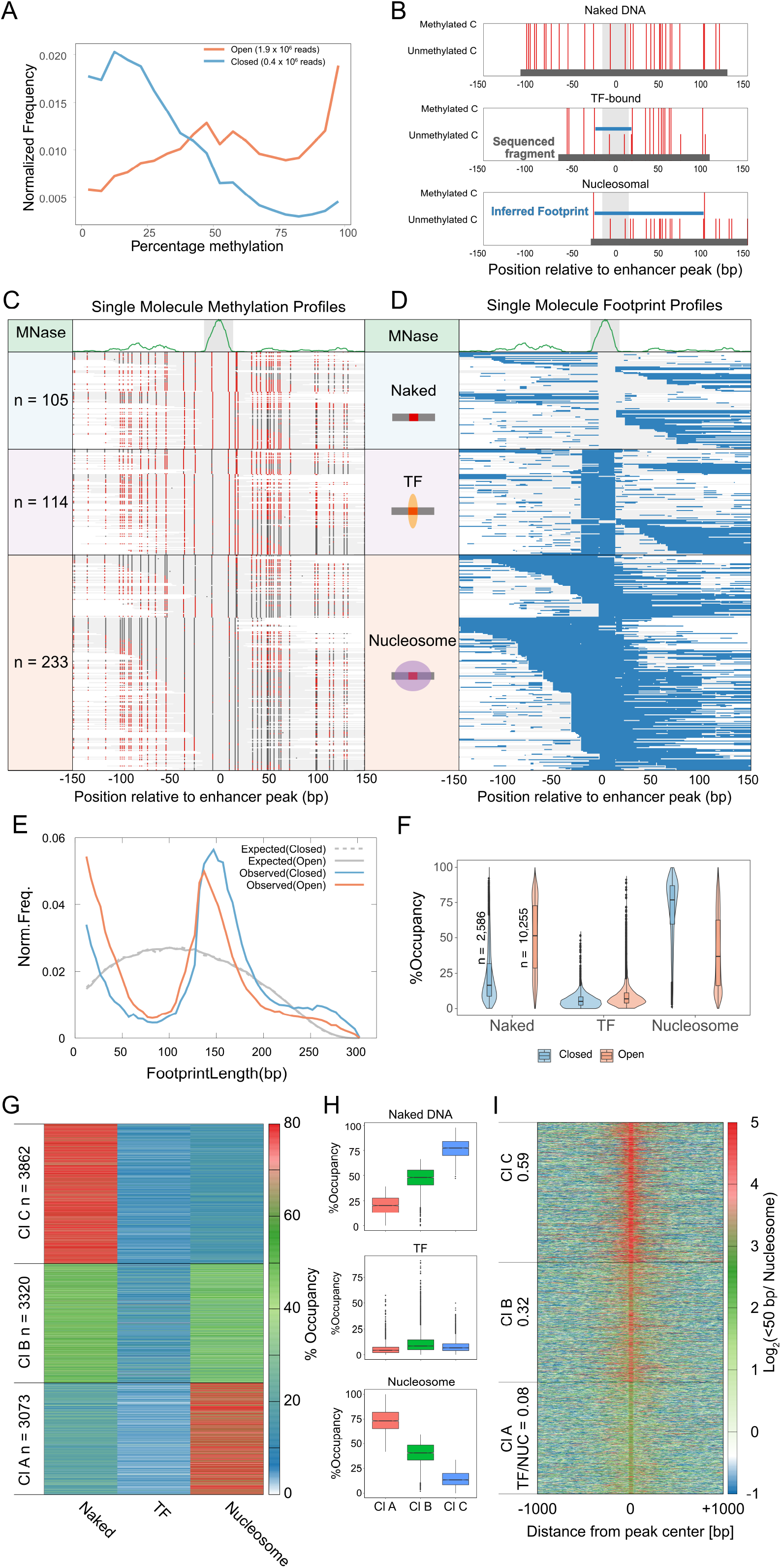
dSMF defines factor binding at enhancers. **A**) Methyltransferase activities around MNase peaks in active and closed enhancers. Percentage methylation defined as the percentage of methylated cytosines (5mC/C) on each read over MNase peaks. **B**) Representative individual bisulfite reads depicting naked (top), TF-bound (middle), and nucleosome-bound (bottom) states of a binding site in an active enhancer (at chr2L:480,305). The footprints called by the algorithm are shown as blue lines and the whole bisulfite read is shown as a dark gray line. **C**) Bisulfite reads mapped to the MNase peak at chr2L:480,305. Unmethylated and methylated cytosines are colored grey and red respectively. **D**) Corresponding reads are plotted in the right panel with called footprints (blue lines). Three clusters (naked, TF-bound, and nucleosome-bound) are shown from top to bottom. **E**) Observed and expected distribution of lengths of footprints defined on bisulfite reads mapping to open and closed enhancers. **F**) The occupancies (percentage of binding states on DNA molecules at active and closed enhancers. **G**) Heatmap of occupancies of three states of each TFBS at open enhancers ordered by the 3 clusters defined using k-means. clusters 1 to 3 are ordered based low to high TF/NUC ratio. **H**) Boxplots of binding state occupancies in the clusters shown in (**G**). **I**) Heatmap of MNase-seq enrichment of short fragments over nucleosomal fragments (Log_2_ (<50 bp/134-160 bp)) plotted relative to primary enhancer peak. Enhancer peaks are plotted in the same order as (**G**).

Methylation reads from dSMF across a representative active enhancer is shown in **Figure 5C**. Based on the size of unmethylated footprints overlapping small protected peaks, we assign one of the three binding states to each read: 1) exposed DNA, with no nucleosomes or bound TFs apparent, 2) a TF-bound structure, where a short unmethylated footprint ≤50 bp in inferred length, or 3) a nucleosome, where the unmethylated footprint is 130-160 bp in length. The proportion of individual reads with each of these states represents the fraction of each state in the population of DNA molecules, and the fraction of time that each structure persists. Thus, for this particular active enhancer, the TF binding site is occluded by a nucleosome 52% of the time, exposed but not bound by a TF 23% of the time, and bound by a TF only 25% of the time. While this does not determine the absolute persistence times of each state, these proportions do imply that the restoration of nucleosomes at this enhancer is relatively slow compared to the on- and off-rates of TF binding.

We then performed analyses of dSMF footprinting across all active enhancers with small protected peaks defined by MNase-seq. Overall, both active and closed enhancers show low frequencies of TF-sized unmethylated footprints, although the group of active enhancers has many more cases of sites with a high proportion of short unmethylated footprints (**Figure 5F**). However, the proportions of exposed DNA and nucleosomal DNA is starkly different between active and closed enhancers. On average, active enhancers are exposed ~50% of the time and occluded by nucleosomes ~30% of the time. In contrast, closed enhancers are exposed only ~20% of the time, and ~75% of molecules are occluded by a nucleosome. Thus, active enhancers are distinguished by extensive eviction of nucleosomes and not high factor occupancy.

In order to probe the diversity of the partitioning of enhancers into the three chromatin structures, we performed k-means clustering across 10,255 small protected peaks defined by MNase-seq. Two of the clusters are characterized by exposed DNA (Cluster C) and nucleosomal structures (Cluster A, **Figure 5G, H**). Cluster B has equivalent proportions of exposed and nucleosomal states and the highest frequency of TF footprinting (**Figure 5G**). When we plotted the occupancy of TF-bound and nucleosome-occluded sites determined by the log_2_ ratio of short fragment enrichment to nucleosome-length fragments in MNase-seq for these clusters, we observed the log_2_ ratio corresponds to the transcription factor/nucleosome ratio defined by dSMF (**Figure 5H**). These MNase-seq ratios in dSMF-defined clusters independently confirm that we are mapping TF binding events at active enhancers. Thus, dSMF both recapitulates protein binding as identified by MNase-seq and enables relative quantification of the exposed state, which is invisible in other methods.

### Cooperative binding is common at active enhancers

We next used dSMF data to analyze active enhancers with primary and secondary small protected peaks, representing regulatory elements with multiple TF binding sites. The median length of DNA molecules in dSMF analysis is 269 bp (**Figure S4**). There are 3 possible binding states of an enhancer with a single small protected peak (exposed, TF-bound, and nucleosome-occluded). Therefore, an enhancer with two small protected peaks has nine potential states, whereas an enhancer with 3 peaks would have 27 states, and so on. To ensure we have sufficient molecules to identify all states, we focused on pairs of two short protected peaks at active enhancers. Out of 11,252 possible pairs of small protected peaks, 5,109 pairs had at least 5X coverage of TF-sized footprints in dSMF data for each peak in the pair. The majority of these pairs have more than 100 reads overlapping both peaks (**Figure 6A**), enabling robust identification of binding states. Indeed, multiple short footprints overlapping two short protected peaks can be readily identified in single dSMF reads (**Figure 6B**), signifying TF co-binding. This read coverage is sufficient to find examples of 3 bound factor footprints in a single read (**Figure 5B**, **bottom**), underlining the potential of dSMF to identify multiple TF binding events in complex regulatory elements.

**Figure 6.**
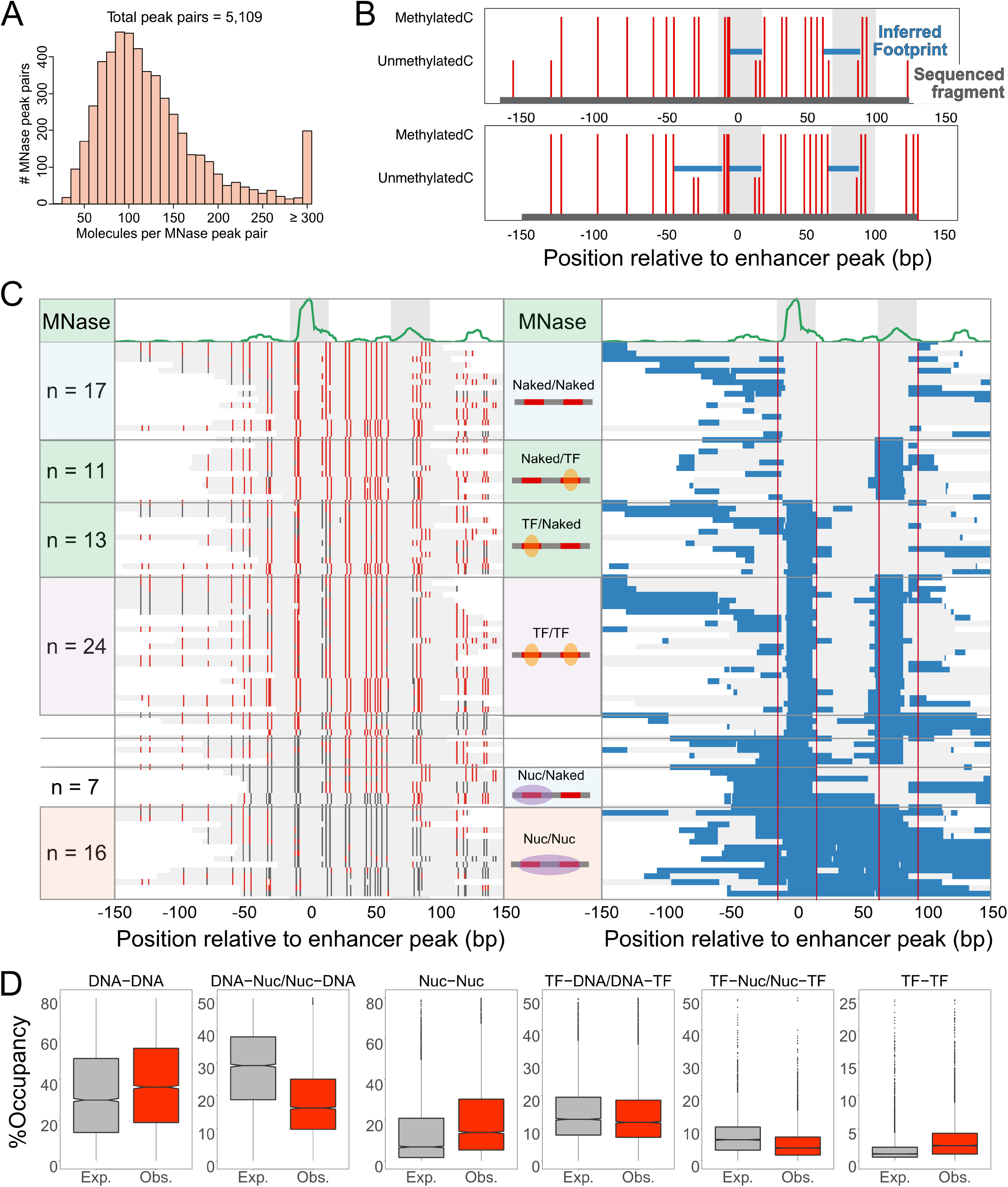
dSMF analysis of TFBS pairs at enhancers. **A**) Distribution of the number of DNA molecules spanning MNase peak pairs used in the co-binding analysis. **B**) Representative individual bisulfite reads depicting two (top), and three (bottom) TF binding events in an intronic active enhancer in the *brat* gene (chr2L:19,155,173). The algorithm-called footprints are shown as green lines and the whole bisulfite read is shown as a purple line. **C**) Bisulfite reads mapped to MNase peak in the *brat* intronic enhancer. Unmethylated and methylated cytosines are colored grey and red respectively. Corresponding reads are plotted in the right panel with called footprints (blue lines). Nine clusters: naked at both sites, naked and TF-bound, TF and naked, TF and TF, TF and nucleosome, nucleosome and TF, Nucleosome and naked, and nucleosome on both sites are shown from top to bottom. The two MNase peaks are 78 bp apart. **D**) Boxplot of the observed and expected prevalence of six states of pairs of short MNase peaks (reduced to six from nine; occupancies of heterogeneous states were merged: for example, occupancies of TF(TFBS1)-Nuc (TFBS2) and Nuc (TFBS1)-TF(TFBS2) were added).

We developed an algorithm to classify binding states at enhancers with two small fragment peaks from methylation profiling (peak pair distance distribution: **Figure S5B**), with additional constraints for co-bound states (see ‘Assigning binding states to DNA molecules’ in Methods). At one such enhancer, we observed reads where one or the other peak site was protected by a TF (one site unmethylated) and reads where both peak sites were protected by TFs (both sites unmethylated), in addition to the exposed and nucleosome-occluded states (**Figure 6C**). We then calculated the prevalence of these states across all enhancers with two peaks (**Figure 6D**). As we observed at elements with single small fragment peaks, the exposed DNA state is the most common (**Figure 6D**). We then calculated the expected probability for each of the states of the peak pairs being independent. Strikingly, homotypic states (DNA-DNA, Nuc-Nuc, TF-TF) have significantly higher observed prevalence than expected (**Figure 6D**), while heterotypic states (DNA-Nuc/Nuc-DNA, TF-DNA/DNA-TF, TF-Nuc/Nuc-TF) are less frequent than expected. Thus, this global analysis of pairs of binding sites within enhancers suggests that elements move in step between co-bound and nucleosome-occluded states.

In groups of enhancers we see cooperativity by MNase-seq. With dSMF, we can score co-binding within individual regulatory elements. We compared observed co-binding frequencies to expected frequencies of co-binding predicted based on methylation state in active enhancers over each small protected peak. We normalized this “cooperativity score” from 0 to 100 (see Methods). The significance of these scores was calculated as a p-value for the observed frequencies of the co-bound state using the hypergeometric test with multiple hypothesis correction to obtain adjusted p-values for all 5,109 peak pairs. We found 31% of peak pairs had a significant (*p*<0.01, median cooperativity score =71) extent of cooperativity (**Figure 7A**). The strength of cooperativity decreases as the distance between small fragment peaks increases, with the largest cooperative effects observed between peaks that are <60 bp apart (**Figure 7C**). More moderate but significant cooperativity also occurs between peaks spaced as far apart as 140 bp, perhaps from indirect cooperativity (Morgunova and Taipale, 2017). In enhancers with at least two peaks, we found that the majority (64%) show at least one cooperative interaction, and 28% of enhancers have more than one such interaction (**Figure 7B**). Thus, cooperative interactions are widespread in regulatory elements across a wide range of spacings that imply multiple mechanisms of synergy.

**Figure 7.**
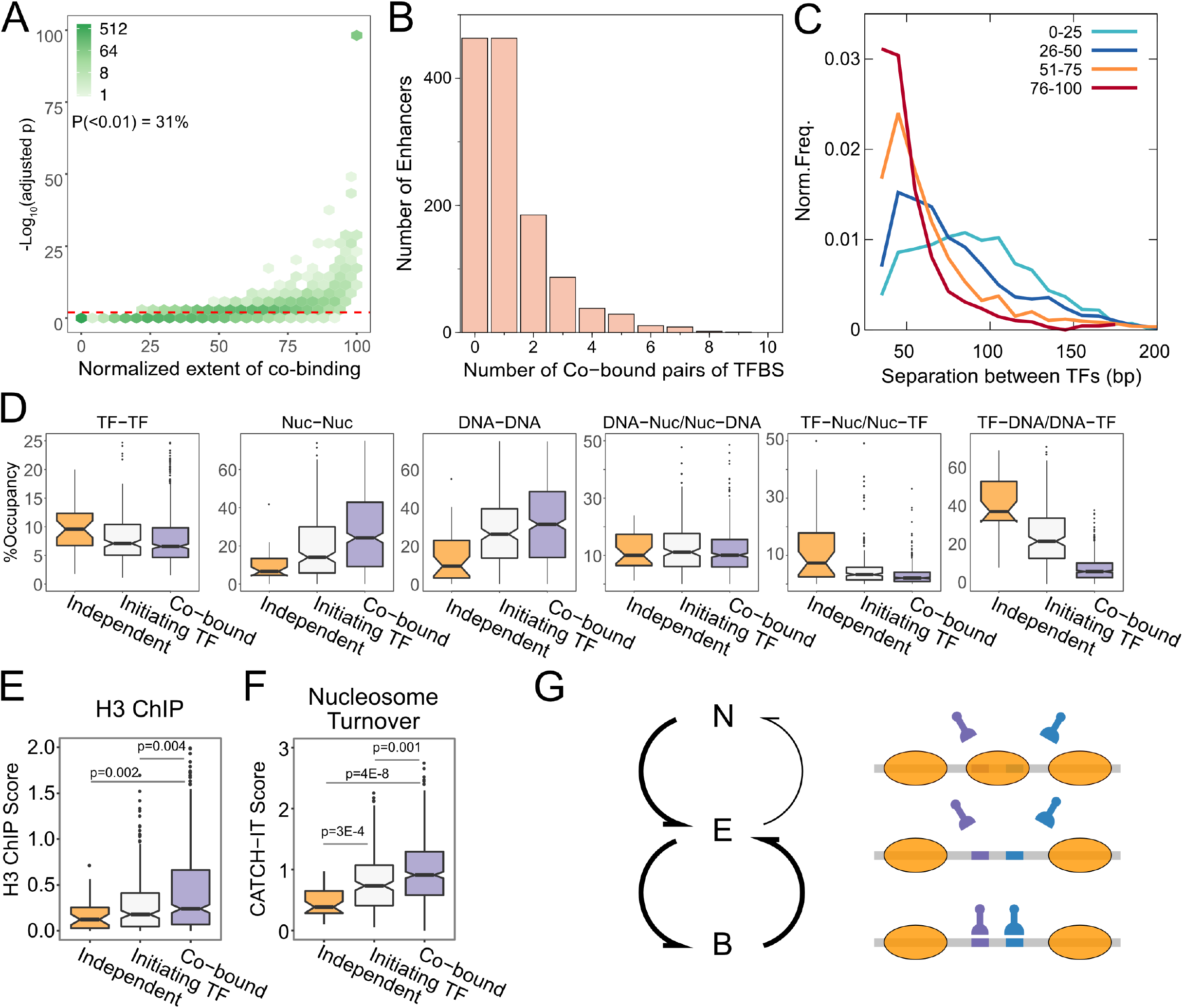
Cooperative binding of transcription factors is characteristic of active enhancers. **A**) Significance level (−log_10_(p-value)) is plotted against normalized extent of co-binding. The p-value is calculated using the hypergeometric test for overlap and p-values were adjusted for multiple hypothesis testing using the Benjamini & Hochberg method. The red dotted line is drawn at p-value = 0.01 (−log_10_(p) = 2). Points above the red line are peak pairs that show significant co-binding. **B**) Distribution of the number of co-binding events observed at enhancers. At about 64% of enhancers, at least one co-binding event is observed. **C**) Distribution of distance between peak pairs plotted for four classes assigned based on their normalized extent of co-binding. **D**) Boxplots of occupancies for six states of binding site pairs plotted for the three classes of cooperative binding defined based on their normalized extent of co-binding and imbalance scores. Independent binders have a non-significant normalized extent of co-binding (p-val>=0.01); Initiating TFs have significant co-binding and an imbalance score ≥2 and co-bound pairs have significant co-binding and imbalance score < 1. **E**) Histone H3 ChIP scores calculated at regions defined by peak pair loci corresponding to three classes described in (**C**). **F**) CATCH-IT nucleosome turnover scores calculated at regions defined by peak pair loci corresponding to three classes described in (**C**). **G**) Schematic of binding state transitions at active enhancers. N refers to nucleosomal, E refers to exposed DNA, and B refers to TF-bound. The thicker arrows represent putative higher rates of transition from nucleosomal to exposed and between exposed and TF-bound states.

In enhancers with two or more small fragment peaks, factors may independently bind on and off from their cognate sites, or binding of one factor may be required to potentiate binding at a second site. To detect such initiating TFs, we compared the frequencies of methylation between peak sites in an enhancer, and if one site had a protection frequency >2-fold of the other site we scored it as a potential initiating site. Together with cooperativity scores, we distinguish three categories: enhancers where two factors appear to bind independently, enhancers where one TF appears to initiate binding of another, and enhancers where cooperativity dominates. Overall, ~30% of enhancers contain an initiating TF site. Thus, the majority of enhancers in the *Drosophila* genome exhibited either binding of cooperative TFs or an initiating TF. We next determined the binding states of these three categories of enhancers. Cooperative enhancers had the highest occupancies of both sites being exposed or nucleosome-bound (**Figure 7D**), indicating that TF cooperativity may effectively displace nucleosomes from high-affinity sequences. On the other hand, independent TF binding occurs at enhancers that appear to be intrinsically nucleosome-depleted (**Figure 7D**). Measures of nucleosome occupancy by ChIP for histone H3 (Mueller et al., 2017) and of nucleosome turnover by metabolic labeling of histones (CATCH-IT) (Teves and Henikoff, 2011) support the idea that enhancers with independent binding of TFs have low nucleosome occupancy and low nucleosome turnover, while enhancers with cooperatively-bound TFs have high occupancy and high turnover of nucleosomes (**Figure 7E, F**). The chromatin dynamics of these enhancers suggests TF cooperativity enables efficient nucleosome displacement at enhancers with high nucleosome affinity, perhaps enabling chromatin regulation of enhancer activity.

## Discussion

Here, we exploit MNase-resistant protections of chromatin to detect bound proteins at high resolution and infer the regulatory architecture of enhancers in the *Drosophila* genome. Enhancers have been thought to have poorly positioned nucleosomes, perhaps corresponding to weak initiation of transcription within elements, but we find that alignment of active enhancers by the factor-protected regions within them resolves chromatin features, revealing that enhancers – like active promoters – are structured and have defined nucleosome-depleted regions. The size and magnitude of NDRs are related to the spacing between factor-protected regions and can span ~400 bp, suggesting that this may be the modular size of enhancer elements. Notably, while factor-protected regions within enhancers often encompass recognizable consensus motifs for known transcription factors, many features and even elements lack any statistically significant motif. As the *Drosophila* transcription factor repertoire has been extensively characterized, this highlights that the rules dictating factor binding *in vivo* remain incomplete. Other aspects of chromatin beyond the sequences directly contacted by transcription factor DNA-binding domains must promote the recognition and effective binding of regulatory sites.

Information guiding factor binding may come from DNA conformation around binding sites (Gordan et al., 2013; Inukai et al., 2017). Additionally, cooperativity between multiple transcription factors in a regulatory element can increase affinity and specificity for weaker consensus motifs (Crocker et al., 2015), for example by dimeric factors (Morgunova and Taipale, 2017; Rastogi et al., 2018; Slattery et al., 2011). Transcription factors juxtaposed on a regulatory element might also enhance the affinity of each factor to DNA *in vitro* (Adams and Workman, 1995; Moyle-Heyrman et al., 2011; Polach and Widom, 1996), but given the fast transient binding of factors *in vivo* (Voss and Hager, 2014; Wilczynski and Furlong, 2010), it has not been clear how widespread factor cooperativity is. We find that while multiple transcription factors do bind independently at some active enhancers, 64% of active enhancers in the fly genome display substantial degrees of factor cooperativity. These cooperative interactions are not due to dimeric factors, since the cases we identify occur between factors that bind regulatory elements that are >30 bp apart. In some cases, cooperativity occurs between factors as far apart as 140 bp. Such long-distance synergies might be due to interacting factors that bridge distant sites, or by effects on nucleosome positioning (Mirny, 2010).

Transcription factor cooperativity correlated with nucleosome occupancy and histone turnover at active enhancers, implying antagonism between transcription factors and histones for DNA. In the context of chromatin, binding of multiple spaced transcription factors competes with nucleosome formation (Mirny, 2010; Moyle-Heyrman et al., 2011; Polach and Widom, 1996). In dynamic nucleosomes where DNA is being unwrapped and rewrapped across the surface of a histone octamer, binding of transcription factors at exposed DNA can block rewrapping of octamers. The efficiency of blocking the restoration of a nucleosome depends on the relative positioning of factor binding sites, where multiple binding sites on one side of a nucleosome are better competitors. Our observation that factor cooperativity occurs predominantly between sites spaced 50 bp apart in active enhancers fits with this idea. In this line of thinking, an important aspect of factor binding site grammars may be loosely constrained arrangements of sites that primarily act to destabilize nucleosomes. As many transcription factors recruit chromatin remodeling enzymes to their binding sites, catalyzed displacement of nucleosomes may also contribute to cooperative occupancy of regulatory elements.

A striking but unexplained observation in single-molecule profiling of eukaryotic chromatin is that factor binding sites are not bound by a cognate factor or occluded in a nucleosome up to ~25% of the time (Krebs et al., 2017; Sönmezer et al., 2020; Stergachis et al., 2020; Vierstra et al., 2020). These observations agree well with single-molecule tracking experiments that show only a small fraction of TFs to be bound stably to chromatin and that most TFs have a short residence time in the order of seconds at stably bound sites (Chen et al., 2014; Paakinaho et al., 2017). How do regulatory elements function when no transcription factor is bound? While transcription factors may often be absent from a regulatory element, part of the answer may lie in that restoration of nucleosomes is slow compared to the binding and release of factors in an active regulatory element (**Figure 7G**). With slow nucleosomal restoration, transient binding of factors maintains a regulatory element in an exposed configuration where factors can cycle on and off. The persistence of histone modifications on flanking nucleosomes may similarly provide a short-term memory of regulatory events when factors are not bound. In these ways, nucleosome dynamics may provide a mechanism to temper stochastic effects of transient factor binding *in vivo*, and thus provide stable regulatory output to direct gene expression.

## Acknowledgements

This work was supported by the RNA Bioscience Initiative, University of Colorado School of Medicine and NIH grants R35GM133434 (S.R.) and R01GM108699 (K.A.). We thank Sujatha Jagannathan and Alexis Zukowski for critical reading of the manuscript.

## Materials and Methods

### Biological materials

Drosophila S2 cells were purchased from Invitrogen and grown to mid-log-phase in HyClone Insect SFX media (GE). We used a primary antibody to Drosophila TRL (Melnikova et al., 2004) (a gift from G Cavalli, Institute of Human Genetics, Université de Montpellier, France) and protein A-micrococcal nuclease fusion (Skene and Henikoff, 2017) (pAMNase, a gift from S. Henikoff, Fred Hutchinson Cancer Research Center, Seattle WA).

### CUT&RUN profiling

We used an immuno-tethered strategy for profiling the binding of the TRL transcription factor in *Drosophila* cells. The CUT&RUN method uses an antibody to a specific chromatin epitope to tether pAMNase at chromosomal binding sites within permeabilized cells (Skene and Henikoff, 2017). The nuclease is activated by the addition of calcium and cleaves DNA around binding sites. Cleaved DNA is isolated and subjected to paired-end Illumina sequencing to map the distribution of the chromatin epitope. CUT&RUN profiling with 1×10^6^ S2 cells and library amplification with 14 cycles of PCR was performed as described (Skene and Henikoff, 2017). Libraries were sequenced for 25 cycles in paired-end mode on the Illumina HiSeq 2500 platform at the Fred Hutchinson Cancer Research Center Genomics Shared Resource. Paired-end reads were mapped to the dm3 version of the D. melanogaster genome (FlyBase.org) using Bowtie2.

### Data and code availability

All datasets were aligned to the dm3 version of the *Drosophila* genome. External datasets used in this study are listed in **Supplementary Table S2**. CUT&RUN profiling for the TRL transcription factors has been deposited in GEO under accession GSEXXXXX and will be made public upon acceptance. The code to reproduce figures is available at https://doi.org/10.5281/zenodo.3979883 (for dSMF analysis). All scripts and pipelines used in the dSMF analysis are available at https://github.com/satyanarayan-rao/protein_binding_at_enhancers.

### Data processing

#### Enhancer identification in S2 cells

We obtained 5,499 STARR-seq summits for Drosophila S2 cells from https://data.starklab.org/publications/yanez-cuna_genomeRes_2014/S2_peakSummits.txt (Yanez-Cuna et al., 2014) and extended the summits by 250 bp on each side.

#### DNase hypersensitivity profiling

Single-end reads from the MODENCODE DNase hypersensitivity dataset were aligned to the dm3 version of the D. melanogaster genome using Novoalign. The fraction of read ends mapped at each nucleotide was multiplied by a constant to give a normalized count at that position.

#### Nucleosome mapping

Nucleosomes carrying the histone H3K27me3 or H3K27ac modifications were identified by selecting 140-220 bp (H3K27me3) or 20-150 bp (H3K27ac) mapped fragments in CUT&RUN profiling (Ramachandran et al., 2017). For profiling nucleosomes and calculating an “H3 ChIP score”, we combined MNase-seq data using 25 U and 100 U of MNase from the T=0 timepoint (control) profiling datasets published by (Mueller et al., 2017), using 140-154 bp mapped fragments. For CATCH-IT profiling, we used 120-174 bp mapped fragments from datasets published by (Teves and Henikoff, 2011). For all nucleosome mapping, the fraction of reads mapped at each nucleotide was multiplied by the size of the Drosophila assembly (139,712,364 bp) to normalize counts at each position, and counts were then aggregated into 10 bp windows.

#### TRL profiling

Binding sites of the TRL transcription factor were mapped using 20-50 bp fragments in ORGANIC native ChIP-seq and CUT&RUN datasets. Coverage was normalized by the size of the *Drosophila* assembly (139,712,364 bp) and counts were aggregated into 10 bp windows.

#### MNase profiling

We used paired-end sequencing data of MNase-digested Drosophila S2 nuclei enriched for small protected fragments by gel-isolation of digested DNA <100 bp (Ramachandran et al., 2017) and aggregated it together with sequenced datasets of total MNase-digested chromatin (Ramachandran et al., 2017; Ramachandran and Henikoff, 2016; Teves and Henikoff, 2011) (**Table S2**). To calculate TF footprint enrichment, we calculated the normalized log2 of TF-protected counts / Nucleosome-protected counts at every basepair in enhancer segments using protected fragments ≤50 bp for TFs and protected fragments 134–160 bp for nucleosomes.

To call TF peaks in enhancers, we normalized counts of centers of ≤50 bp protected fragments in aggregated MNase-seq data at basepair-resolution, and then smoothed counts with a 5 bp sliding window, generating the “≤50 high-resolution track”. We then intersected this track with STARR-seq enhancers (Yanez-Cuna et al., 2014) and called peaks using a custom script (deposited at https://github.com/srinivasramachandran/Dm-Enhancer-MNase). Each active enhancer was centered at its tallest peak, and Z-scores were calculated at the centers of ≤50 bp fragments ±200 bp from the central peak, and k-means clustering (k=9) was performed over the Z-score matrix.

For identifying motifs underlying small protected peaks, we used FIMO (Grant et al., 2011) with default parameters on the peaks called from ≤50 bp protection track with consensus motifs from Fly Factor Survey (Enuameh et al., 2013) and JASPAR (Fornes et al., 2020). Recovered motifs were then filtered for the corresponding TFs that are expressed in S2 cells with expression scores ≥10 in S2-DRSC microarray profiling data or ≥300 RPKM in S2-DRSC RNA-seq profiling data (Cherbas et al., 2011).

For V-plots of clustered enhancers, we binned small protected fragments by their fragment lengths and positions relative to centered enhancer summits. The fragment length vs. fragment midpoint position 2D histogram was normalized by the number of nucleosomal fragments (142-152 bp in length) mapping ±15 bp from the enhancer peak to plot the enrichment of fragments over nucleosome density at the peak.

For V-plot simulations, we first defined protections that corresponded to TF binding. We set the probability of fragment ends at the edge of protections as 0.01. We set the probability of fragment ends at the rest of the region of interest (−270 to 270 bp from central protection) as 0.0018. The probability of fragment ends within the protections was set to 0. We then simulated 2×10^6^ fragments of lengths 25 to 125 bp based on the probability of fragment ends at each location in the region of interest. We then calculated the density of fragment length vs. fragment midpoints. We normalized the density based on each cluster’s length distribution. Thus, each cluster’s simulated V-plots reproduced the probability of fragment ends and length distribution of that cluster. We generated a total of 2×10^6^ fragments from 3 states: TF bound only at the central peak, TF bound only at the secondary peak, and finally, TFs bound at both central and secondary peaks. We fractionated the 3 states to represent different levels of cooperativity, where cooperativity is defined by the following ratio:

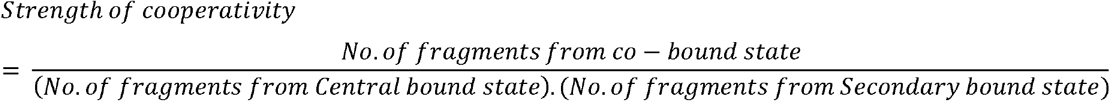

We generated 100 V-plots for each value of strength of cooperativity.

#### dSMF sequence alignment

We used dual-enzyme single-molecule footprinting dSMF profiling data of Drosophila S2 cells published by (Krebs et al., 2017). We downloaded 150bp paired-end reads and used Trim-Galore (https://github.com/FelixKrueger/TrimGalore) to remove sequencing adapters and Bismark (Krueger and Andrews, 2011) to align bisulfite sequences to the dm3 assembly. Biological and technical replicates were merged to create a single alignment file for the downstream analysis. DNA methylation calls on non-CpG/GpC dinucleotides (~10% of the total) were discarded.

#### DNA molecule preparation and footprint calls

To infer protein binding events at single DNA molecule resolution, we examined only overlapping (≥0 bp) aligned reads and assessed the methylation of single DNA molecules. We called footprints of DNA regions protected by chromatin proteins if at least one cytosine was unmethylated between methylated cytosines in a read. Footprints <10 bp in length were discarded, as transcription factors typically protect ~10 bp or more. Special consideration was mapped reads with unmethylated cytosines but devoid of footprints; these represent a nucleosome bound at either of the edges of the read. Such nucleosomal footprints are called if the length of a DNA segment with unmethylated cytosines on edges was >130 bp. Footprints that were separated by one bp (“wobble”) were merged to define a longer footprint. To estimate potential footprint length profiles on reads as a baseline for observed footprints, we defined potential footprints by taking all combinations of three cytosines (^n^C_3_) in mapped reads.

#### Assigning binding states to single factor sites in dSMF reads

We defined a ±15 bp window around each small fragment peak center within active STARR-seq enhancers as the putative TF binding site. Any dSMF read with all cytosines methylated across the segment or with unmethylated cytosines limited to <10 bp was annotated as an exposed (E) binding site. An unmethylated footprint spanning 10-50 bp was annotated as a TF-bound (T) site, and unmethylated footprints >130 bp were annotated as nucleosome-occluded (N) sites. In cases of more than one unmethylated footprint in a read, the footprint with the largest overlap to the TF peak was assigned as its chromatin structure. Footprints at the edges of a read are special cases that were only retained and annotated as nucleosome-occluded if the edge unmethylated footprint spanning a binding site was >50 bp in length, *ie.* greater than expected for protection by a TF and consistent with nucleosomal protection.

#### Assigning chromatin structures to pairs of binding sites in dSMF reads

Three possible structures at a TF binding site gives nine possible structures for pairs of binding sites. We used the above criteria for structure annotation, with the additional defined structures: reads with an unmethylated footprint <100 bp encompassing both binding sites were annotated as 2 co-bound (C) TFs (**Figure S5A**).

We calculated the normalized extent of transcription factor co-binding as the ratio of two probability terms using the following formulae

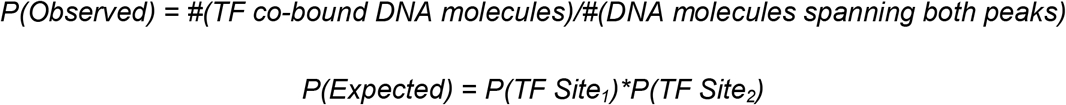

Where P(TF Site_i_) = #(TF-bound DNA molecules at Site_i_)/#(DNA molecules spanning both peaks)

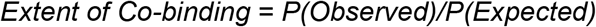

It can be seen that the extent of the co-binding term is directly proportional to the total number of DNA molecules spanning both sites. To remove this bias, we multiply by the extent of co-binding by the total TF binding fraction:

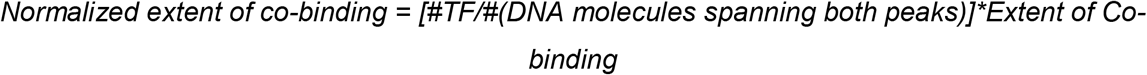

Figure S6 depicts the effect of normalization. We use this normalized extent of co-binding values throughout analysis.

To infer preferred binding sites within an enhancer, we calculate an Imbalance Score using the following formulae:

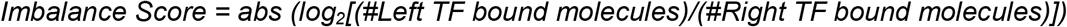

Thus, more TF-bound molecules at one site compared to another gives a high imbalance score. We annotated a binding site in a pair of sites as an Initiating site if it had an imbalance score ≥1.

---

**Figure S1. TRL binding at active enhancers. A**) Heatmaps comparing enrichment of MNase short fragments and TRL ORGANIC plotted relative to the primary peak of MNase short protection at enhancers with no TRL motifs. **B**) Schematic showing the definition of primary and secondary sites based on the presence of TRL motifs and the expected outcomes from MNase, ORGANIC, and CUT&RUN in the context of TRL binding at primary or secondary sites. Two cases are defined based on TRL binding. Case 1: where TRL (shown as green eclipse) is binding at the primary peak of the enhancer (the left extreme of the DNA) and non-TRL TF binds at the secondary peak; and Case 2: the opposite of case 1. MNase and CUT&RUN are expected to release both bound TRL and other TFs, but ORGANIC will only recover bound TRL.

**Figure S2. V-plots at enhancer clusters.** Fragment midpoint versus fragment length plot (V-plot) centered at the primary peak for enhancers for Clusters 1 (**A**), 2 (**B**), 7 (**C**), 8 (**D**), and 9 (**E**). Fragments 100-200 bp (top) and 0-100 bp (bottom) are plotted separately because <100 bp fragments are over-represented due to experimental size selection for enriching TF protections.

**Figure S3. Distribution of co-bound species relative to primary enhancer peak. A**) The average density of centers of 40±5 bp fragments calculated from the V-plot is plotted relative to the primary enhancer peak for Cluster 3. **B**) The average density of 40±5 bp and 82±5 bp fragments calculated from the V-plot of Cluster 4. The gray line represents the density of 82±5 bp fragments calculated from Cluster 3. **C**) The average density of 40±5 bp and 100±5 bp fragments calculated from the V-plot of Cluster 5. The gray line represents the density of 100±5 bp fragments calculated from Cluster 3. **D**) The average density of 40±5 bp and 122±5 bp fragments calculated from the V-plot of Cluster 6. The gray line represents the density of 122±5 bp fragments calculated from Cluster 3.

**Figure S4: Distribution of DNA molecule lengths used in dSMF analysis.** Distribution of DNA molecules mapped to all MNase peaks from active and closed enhancers. A DNA molecule indicates overlapping/adjacent read mate pairs from bisulfite sequencing reads.

**Figure S5: DNA molecule selection criteria for co-binding analysis. A**) Top left: Methylation vector plot for DNA molecules example peak shown in Figure 5C. These molecules are prepared using paired-end reads with sam flag 99 and 147. Red dot: methylated cytosine; Black dot: unmethylated cytosine; Purple dot: Fill-in; D: Naked DNA state; N: Nucleosome state; T: TF-bound state; X: molecule to be discarded. Top right: footprint called using methylation patterns (5mC-(C)_n_-5mC); orange dot: footprint. Here only one read (labeled D-X in right) is discarded as it can be seen that the footprint called for the right MNase peak corresponds to the right edge of the DNA molecule, thus not considered in the analysis. Bottom left: Same representation for DNA molecules prepared from paired-end reads with sam flag 163 and 83. Bottom right: Footprint representation. Three DNA molecules are discarded from analysis (two with X-T and one with X-N). Please see the interpretation of sam flags here: https://broadinstitute.github.io/picard/explain-flags.html **B**) Distribution of distance between MNase peak pairs. Total MNase peak pairs (n = 5,109) with at least 5 TF-bound DNA molecules for both TFBSs.

**Figure S6: Correcting for bias in the extent of co-binding calculation.** Left: before normalization. Low percentage TF (total TF bound molecules)/(Total molecules) lead to an unreliably high extent of co-binding. Right: Corrected for the bias (see Methods).

**Table S1. List of motifs identified at short MNase peaks in active enhancers**

**Table S2. External datasets used in this study**

## Notes

### Competing Interest Statement

The authors have declared no competing interest.

