## Supplementary material for "Cooperative Binding of Transcription Factors is a Hallmark of Active Enhancers": Table S1

**Table S1. List of motifs identified at short Mnase peaks in active enhancers**

| | N | Percentage |
|---|---|---|
| Total Peaks | 16542 | 100.0 |

| | | |
|---|---|---|
| No Motif | 6021 | 36.4 |

| Motif | N | Percentage |
|---|---|---|
| Trl | 1280 | 7.7 |
| lola | 1124 | 6.8 |
| Mad | 531 | 3.2 |
| da | 526 | 3.2 |
| Aef1 | 407 | 2.5 |
| Adf1 | 294 | 1.8 |
| crol | 287 | 1.7 |
| br | 225 | 1.4 |
| CG3065 | 215 | 1.3 |
| pho | 212 | 1.3 |
| kay | 206 | 1.2 |
| CG4360 | 204 | 1.2 |
| dl | 175 | 1.1 |
| tgo | 173 | 1.0 |
| jim | 170 | 1.0 |
| EcR | 146 | 0.9 |
| ttk | 142 | 0.9 |
| pnr | 138 | 0.8 |
| pan | 137 | 0.8 |
| M1BP | 135 | 0.8 |
| h | 131 | 0.8 |
| tai | 131 | 0.8 |
| Med | 130 | 0.8 |
| z | 125 | 0.8 |
| Dref | 118 | 0.7 |
| CG5953 | 109 | 0.7 |
| cnc | 101 | 0.6 |
| her | 99 | 0.6 |
| srp | 98 | 0.6 |
| crp | 97 | 0.6 |
| CTCF | 93 | 0.6 |
| CG12236 | 92 | 0.6 |
| luna | 92 | 0.6 |
| Bgb | 91 | 0.6 |
| GATAd | 89 | 0.5 |
| CG11504 | 86 | 0.5 |
| schlank | 83 | 0.5 |

| | | |
|---|---|---|
| Su(H) | 81 | 0.5 |
| ZIPIC | 78 | 0.5 |
| aop | 76 | 0.5 |
| exd | 73 | 0.4 |
| ftz-f1 | 71 | 0.4 |
| HLH4C | 71 | 0.4 |
| Cf2 | 68 | 0.4 |
| Sox14 | 67 | 0.4 |
| Hsf | 66 | 0.4 |
| ken | 65 | 0.4 |
| sd | 64 | 0.4 |
| cic | 63 | 0.4 |
| pfk | 59 | 0.4 |
| CG12155 | 58 | 0.4 |
| Ets97D | 55 | 0.3 |
| foxo | 55 | 0.3 |
| CG8765 | 51 | 0.3 |
| Irbp18 | 47 | 0.3 |
| phol | 47 | 0.3 |
| Stat92E | 47 | 0.3 |
| mamo | 42 | 0.3 |
| bowl | 39 | 0.2 |
| Zif | 39 | 0.2 |
| CrebA | 37 | 0.2 |
| brwl | 36 | 0.2 |
| Hnf4 | 33 | 0.2 |
| su(Hw) | 33 | 0.2 |
| chinmo | 32 | 0.2 |
| pnt | 32 | 0.2 |
| tgo_sima | 31 | 0.2 |
| ct | 30 | 0.2 |
| CG5180 | 26 | 0.2 |
| D19B | 26 | 0.2 |
| Rel | 26 | 0.2 |
| NFAT | 25 | 0.2 |
| bigmax | 23 | 0.1 |
| CG33557 | 22 | 0.1 |
| shn | 22 | 0.1 |
| Eip75B | 21 | 0.1 |
| cwo | 20 | 0.1 |
| Mes2 | 20 | 0.1 |
| NK7.1 | 20 | 0.1 |
| Coop | 17 | 0.1 |
| jigr1 | 16 | 0.1 |
| cyc | 15 | 0.1 |

| | | |
|---|---|---|
| Pdp1 | 15 | 0.1 |
| CG12768 | 13 | 0.1 |
| Mnt | 11 | 0.1 |
| Max | 10 | 0.1 |
| fru | 9 | 0.1 |
| CG6276 | 8 | 0.05 |
| Met | 7 | 0.04 |
| Hr78 | 6 | 0.04 |
| BEAF-32 | 5 | 0.03 |
