## Supplementary material for "Cooperative Binding of Transcription Factors is a Hallmark of Active Enhancers": Table S2

| **S. No.** | **Dataset** | **Figures** | **GEO Accession ID** | **Reference** |
| --- | --- | --- | --- | --- |
| 1 | Short and Long Fragments Sequenced after MNase | Figure 1, 2, 3, 4, and 7 | GSM1974516 | (Ramachandran and Henikoff, 2016) |
| 2 | Short and Long Fragments Sequenced after MNase | Figure 1, 2, 3, 4, and 7 | GSM1974518 | (Ramachandran and Henikoff, 2016) |
| 3 | Short and Long Fragments Sequenced after MNase | Figure 1, 2, 3, 4, and 7 | GSM1974522 | (Ramachandran and Henikoff, 2016) |
| 4 | Short and Long Fragments Sequenced after MNase | Figure 1, 2, 3, 4, and 7 | GSM1974524 | (Ramachandran and Henikoff, 2016) |
| 5 | Short and Long Fragments Sequenced after MNase | Figure 1, 2, 3, 4, and 7 | GSM1974526 | (Ramachandran and Henikoff, 2016) |
| 6 | Short and Long Fragments Sequenced after MNase | Figure 1, 2, 3, 4, and 7 | GSM2592578 | (Ramachandran et al., 2017) |
| 7 | Short and Long Fragments Sequenced after MNase | Figure 1, 2, 3, 4, and 7 | GSM2592579 | (Ramachandran et al., 2017) |
| 8 | Short and Long Fragments Sequenced after MNase | Figure 1, 2, 3, 4, and 7 | GSM763030 | (Teves and Henikoff, 2011) |
| 9 | H3K27me3 CUT&RUN | Figure 1 | GSM3424765 | (Ahmad and Spens, 2019) |
| 10 | H3K27ac CUT&RUN | Figure 1 |  | This study |
| 11 | DNase-seq | Figure 1 | MODENCODE 3324 | [Source](https://www.encodeproject.org/experiments/ENCSR834VXA/) |
| 12 | Trl Native ChIP-seq | Figure 2, 7 | GSM1111722 | (Kasinathan et al., 2014) |
|  | Trl Native ChIP-seq | Figure 2, 7 | GSM1111723 | (Kasinathan et al., 2014) |
| 13 | H3 ChIP-seq | Figure 6 | GSM2521698 | (Mueller et al., 2017) |
| 14 | H3 ChIP-seq | Figure 6 | GSM2521699 | (Mueller et al., 2017) |
| 15 | H3 ChIP-seq | Figure 6 | GSM2521702 | (Mueller et al., 2017) |
| 16 | H3 ChIP-seq | Figure 6 | GSM2521703 | (Mueller et al., 2017) |
| 17 | CATCH-IT | Figure 6 | GSM763034 | (Teves and Henikoff, 2011) |
| 18 | Trl CUT&RUN | Figure 7 |  | This study |
| 19 | Bisulfite sequencing data used in dSMF analysis | Figure 4, 5 and 6 | GSM2050819 | (Krebs et al., 2017) |
| 20 | Bisulfite sequencing data used in dSMF analysis | Figure 4, 5 and 6 | GSM2050820 | (Krebs et al., 2017) |
| 21 | Bisulfite sequencing data used in dSMF analysis | Figure 4, 5 and 6 | GSM2050821 | (Krebs et al., 2017) |
| 22 | Bisulfite sequencing data used in dSMF analysis | Figure 4, 5 and 6 | GSM2050822 | (Krebs et al., 2017) |
