## Supplementary figures and images for "Cooperative Binding of Transcription Factors is a Hallmark of Active Enhancers"

### Figure S1

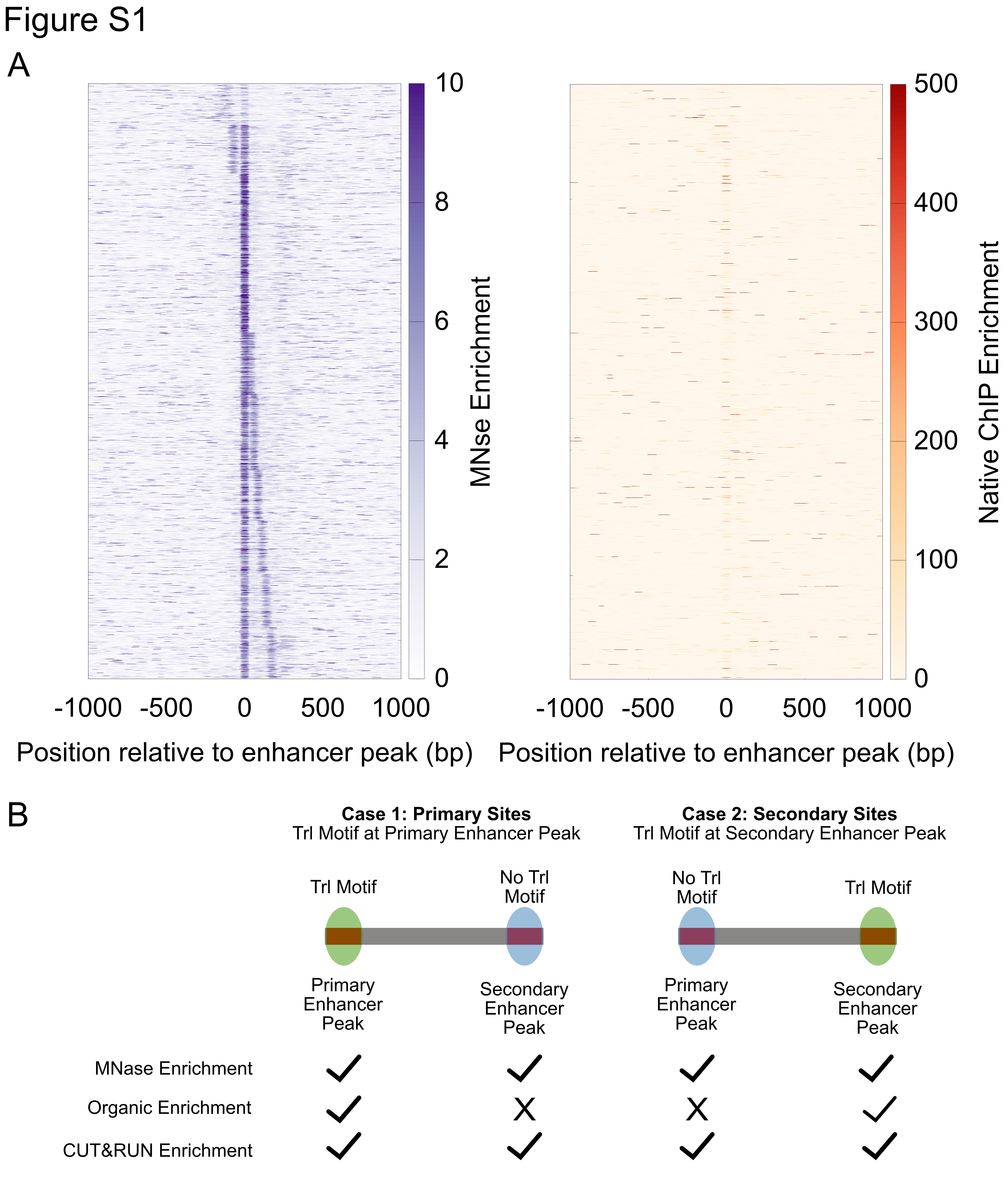

### Figure S2

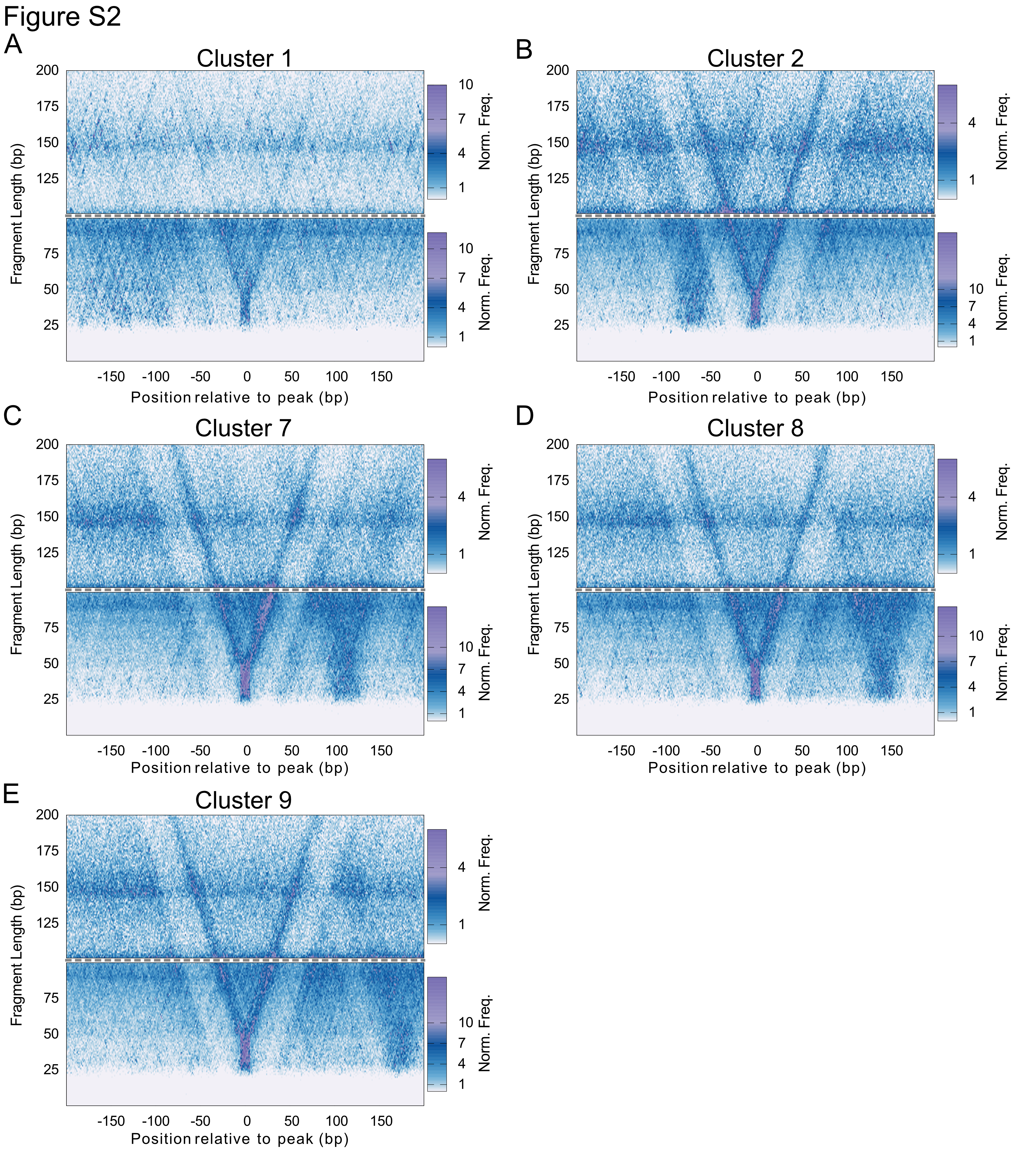

### Figure S3

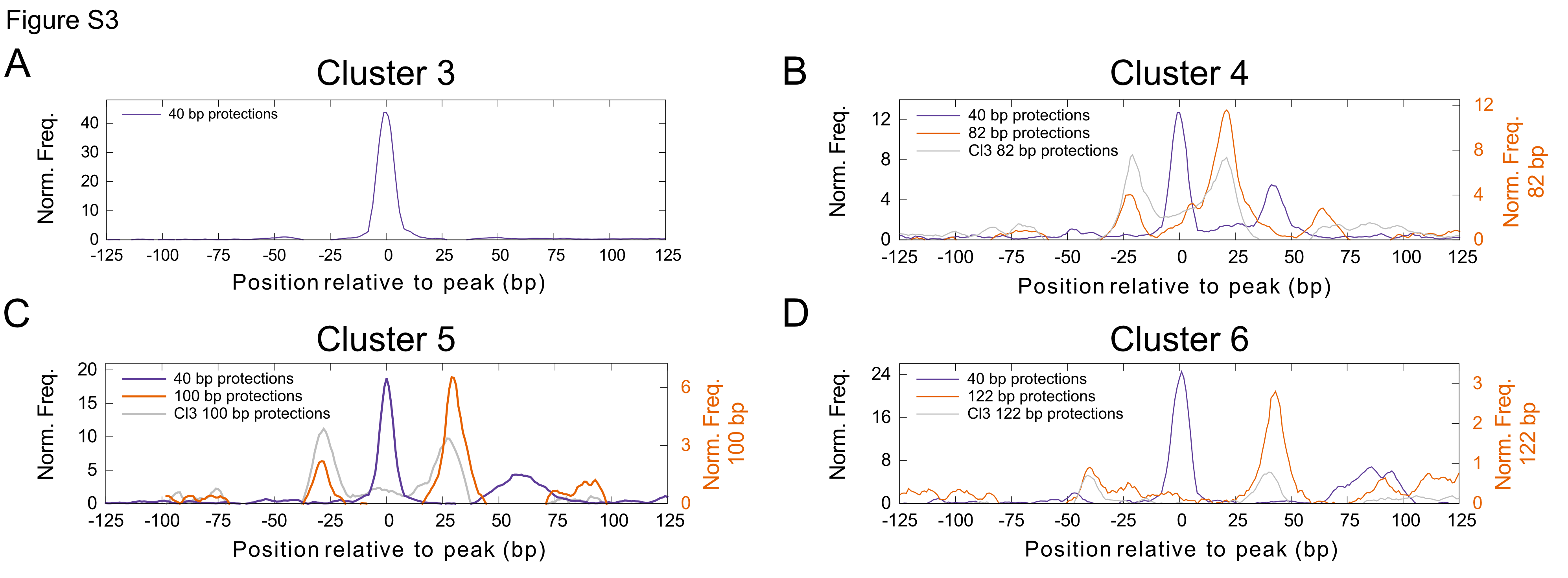

### Figure S4

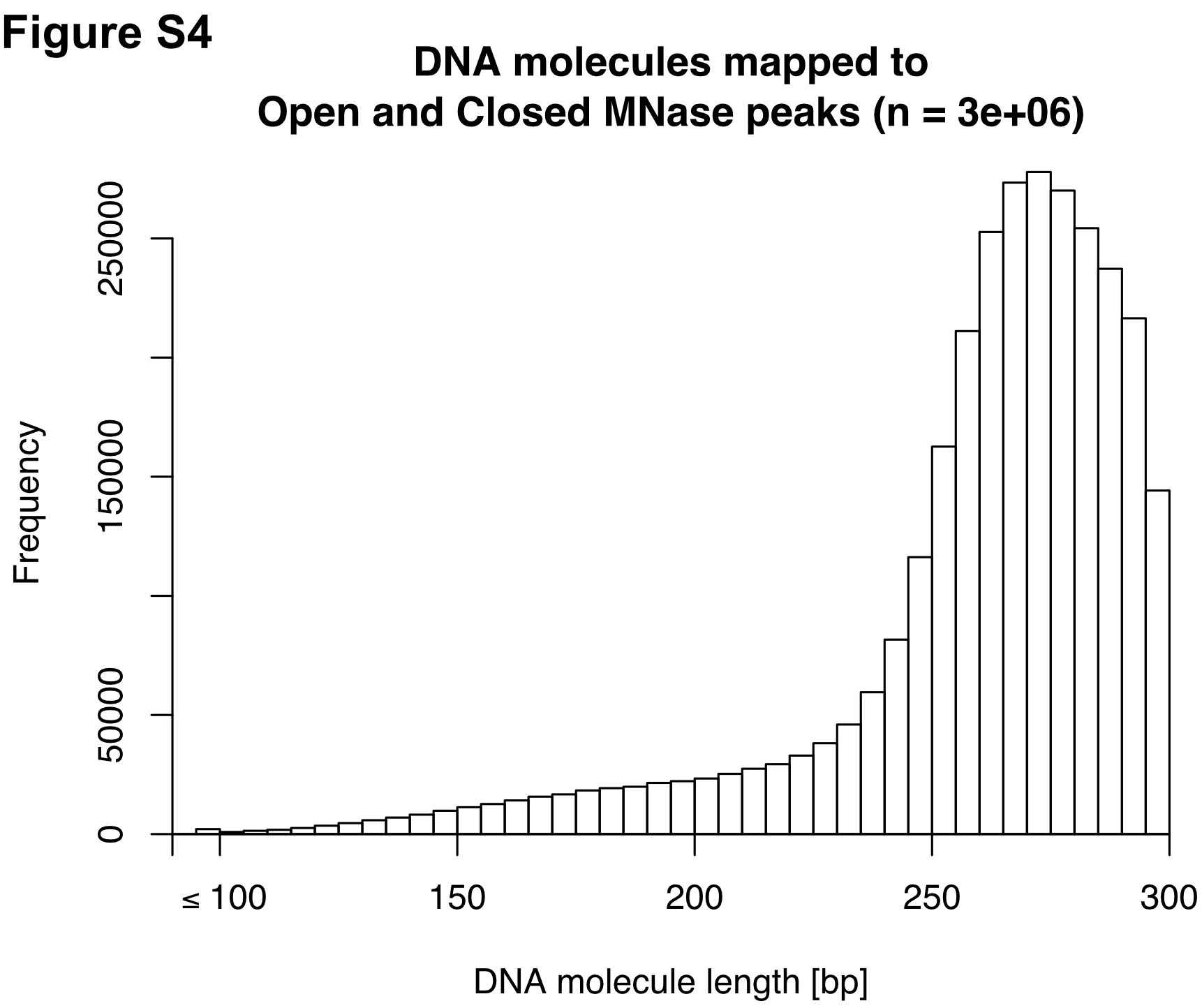

### Figure S5

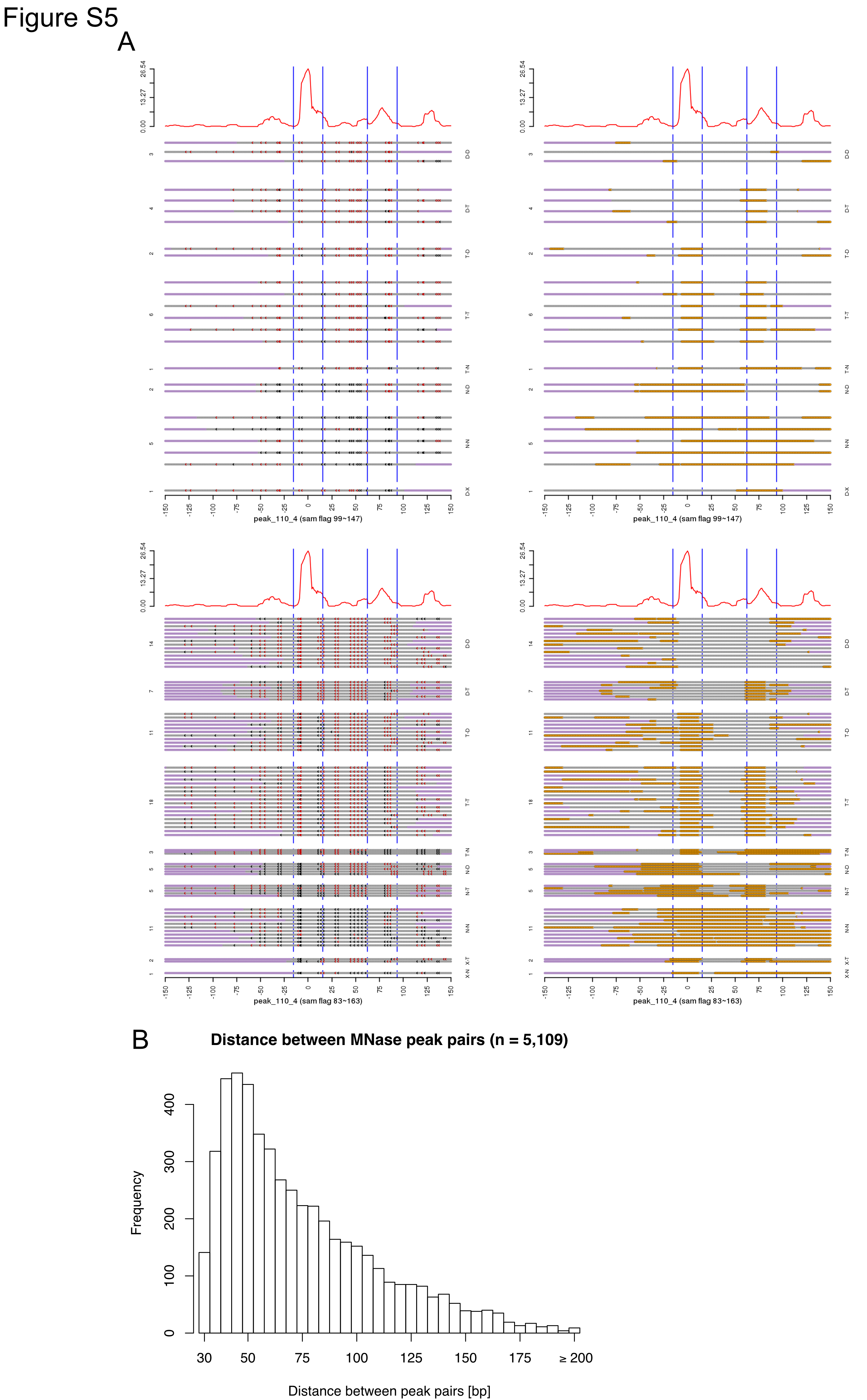

### Figure S6

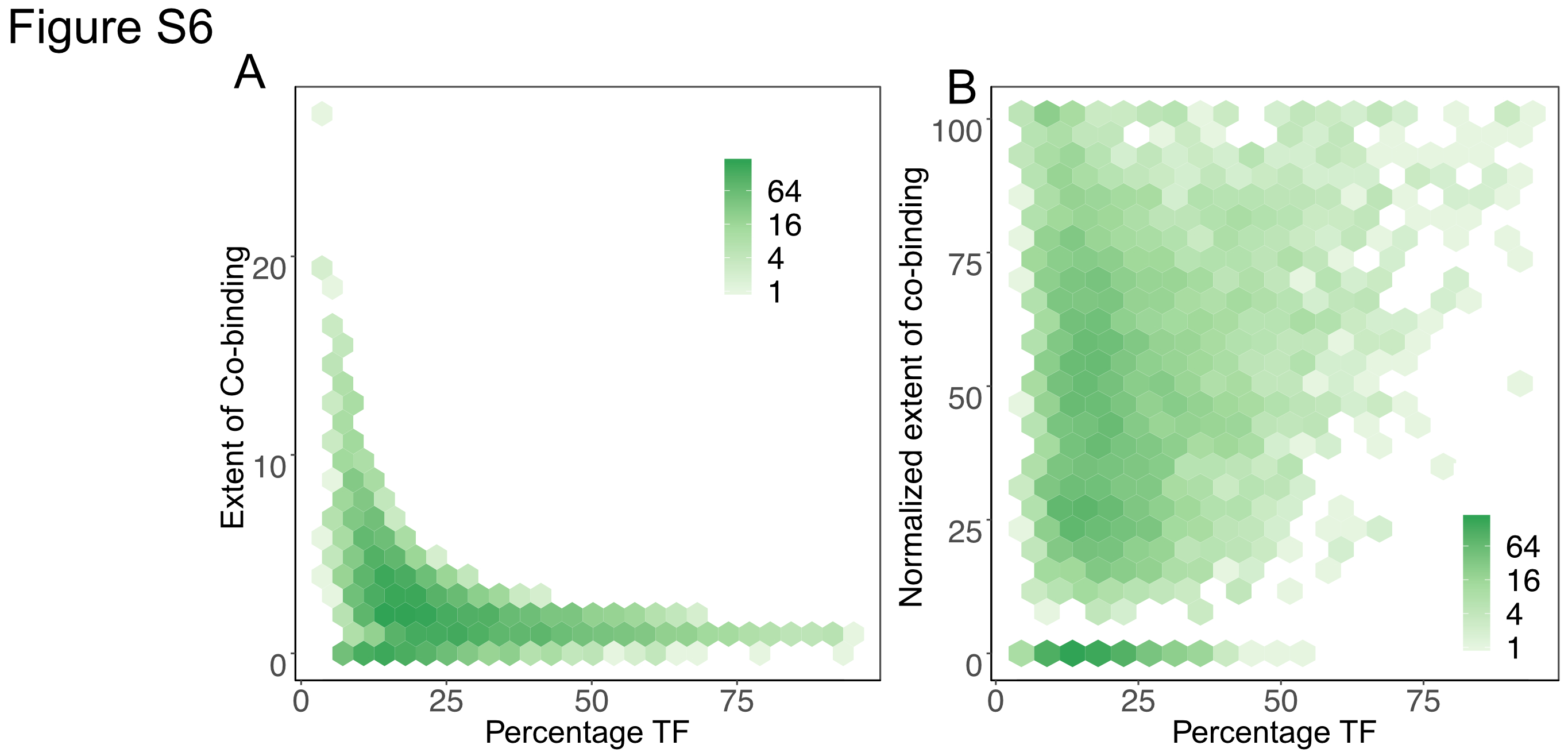
